# FRET-guided integrative modeling resolves Mg^2+^-dependent RNA tertiary-contact formation

**DOI:** 10.64898/2026.09.17.752353

**Authors:** Felix Erichson, Mirko Weber, Vanessa Schumann, Mara Henschel, Josephine Meitzner, Fabio D. Steffen, Patrick K. Quoika, Richard Börner

## Abstract

Tertiary contact interactions between GNRA tetraloops and their receptors are recurrent motifs that stabilize complex RNA folds and in the context of ribosome biogenesis contribute to ribosomal subunit maturation. High-resolution structure determination methods typically yield single structural snapshots, which may not fully describe the conformational collection populated by such contacts in solution. Here, we investigate a ribosomal RNA tertiary contact formed by a GAAA tetraloop and a kissing loop acting as its receptor using single-molecule FRET-guided integrative modeling. We used a minimal RNA model construct in which the tetraloop and its kissing loop receptor are connected by flexible poly(A)-linkers, assembled its bound state from the native ribosomal cryo-EM geometry, and combined restrained and unrestrained all-atom molecular dynamics simulations with *in silico* FRET and dynamic fluorescence anisotropy predictions to compare with the respective experiments under varying salt conditions. The modeled bound state matches the smFRET distribution only at non-physiologically high Mg^2+^ concentrations. At 10 mM Mg^2+^, however, the single-molecule FRET distribution remains heterogeneous and is reproduced by mixtures of bound and unbound conformations, indicating their coexistence under intermediate Mg^2+^ conditions. Thus, smFRET-guided integrative modeling resolves subpopulations invisible to static high-resolution structures and ensemble FRET, providing a route to validate and refine RNA structural models and conformational ensembles in solution.

**TOC-Figure:** 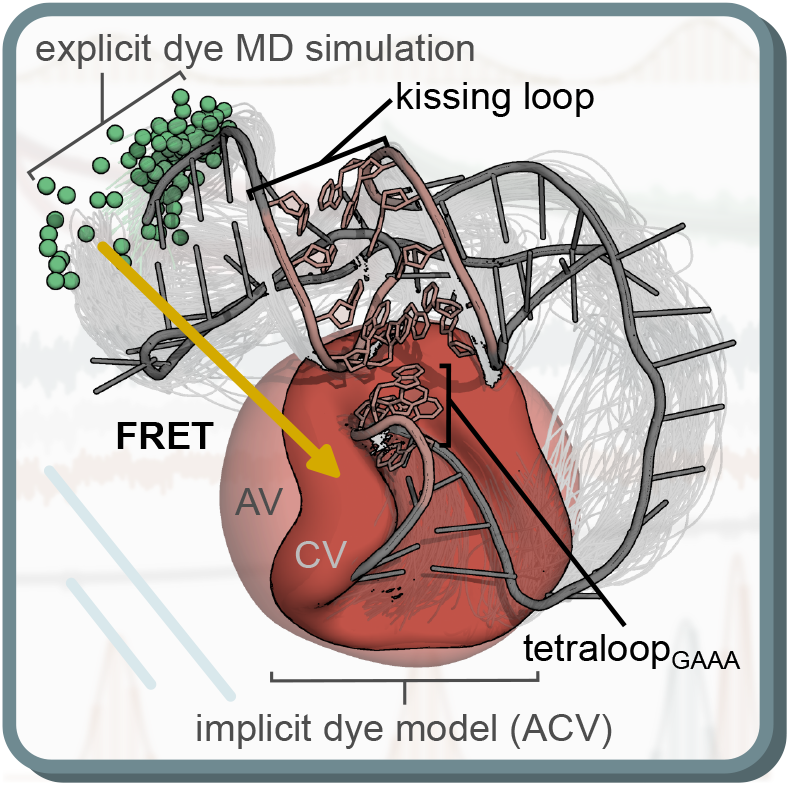

## Introduction

RNA molecules are central to the assembly of complex cellular machines such as the ribosome, where proper folding is essential for the formation of a fully mature and functional ribosomal complex [1]. In this way, RNA folding proceeds as a quasi-hierarchical process, involving a network of tertiary RNA–RNA contacts, stabilizing the three-dimensional fold throughout consecutive steps of folding intermediates and rearrangements [2–6]. However, formation of such tertiary interactions represents a challenge, as they are flexible and arise from sequentially and spatially separated secondary-structure elements, resulting in a vast conformational space [7–9]. Chaperone-like RNA-binding proteins and divalent metal ions, such as Mg^2+^, help to constrain and stabilize the RNA structure beyond canonical base pairing [9–14].

One ubiquitous ribosomal RNA tertiary contact is the tetraloop–receptor (TLR) interaction, in which an A-minor motif engages its receptor domain during ribosomal maturation [4, 15, 16]. In yeast rRNA, this contact is formed between a GAAA tetraloop (TL_GAAA_) and a kissing loop (KL), the latter acting as the TLR and critically shaping the dynamics of large molecular machineries [17, 18]. This tertiary contact was previously investigated by ensemble FRET measurements using a minimal RNA model construct that comprises the H22/H88 KL and the H68 TL_GAAA_ connected by single-stranded poly(A)-linkers [14]. Characterizing this construct *in vitro* with and without the RNA binding protein Puf6 in a range of Mg^2+^ concentrations at room temperature showed that the compact, high-FRET state is reached by increasing the concentration of Mg ^2+^ or, in the absence of Mg^2+^, by Puf6 alone. The latter thereby mimics the stabilizing role of divalent metal ions on the bound conformation. The construct therefore provides a well-defined experimental system in which the formation of a long-range ribosomal RNA tertiary contact can be followed as a function of the surrounding ionic conditions.

Atomic-resolution insights of RNA structures are experimentally achieved by means of X-ray crystallography [19], NMR spectroscopy [20], and cryo-EM [21], with crystallography still accounting for the majority of experimentally resolved RNA structures [22]. However, these methods yield single snapshots or a limited set of final structures, whereas RNA folding and function rely on transitions between multiple, inherently dynamic conformational states [23–25]. In contrast, complementary solution-based techniques such as double electron–electron resonance (DEER) spectroscopy [26, 27] and Förster resonance energy transfer (FRET) [28] probe the dynamic rearrangement at the nanometer scale with local distance trajectories of site-specifically introduced labels. Here, single-molecule FRET (smFRET) observes single molecule dynamics on timescales ranging from nanoseconds to hours [28], so that coexisting subpopulations and their interconversion along a folding trajectory become visible.

However, these so called low-resolution techniques will not provide the all-atom representation underlying the observed dynamic conformational RNA structure ensemble. Instead, RNA 3D structure prediction tools such as RNAComposer [29, 30], Rosetta FARFAR2 [31–33], and AlphaFold3 [34] reliably reproduce secondary-structure elements and local motifs, eventually yielding RNA 3D structure collections [35]. Although remarkable progress has been made to give structure representation over a variety of secondary and tertiary structure elements, long-range tertiary contacts, in particular non-canonical interactions and pseudoknot-like topologies, remain challenging in RNA 3D structure prediction [36–38]. Knowledge-based 3D structure assembly provides a complementary route, as individual elements from experimentally resolved full structures can be combined into a cohesive RNA 3D model in which the tertiary arrangement is imposed explicitly. While the individual building blocks are experimentally supported, their combination into one construct is not. Thus, the assembled model remains a hypothesis and needs to be tested against experimental observables [38] using well established integrative modeling approaches [39].

Although structure prediction not only yield single static conformations, but a collection of structures [35], potential dynamics or transitions between those states are not covered. Molecular dynamics (MD) simulations, however, add this dynamic dimension by sampling the conformational space around a given structure, either freely or restrained [40–42]. On the nanosecond-to-microsecond timescale, which is nowadays accessible in MD simulations, fast local rearrangements and the stability of a given conformation can be captured, while slower transitions between folding states remain mostly computationally out of reach [43, 44]. FRET-guided integrative modeling combines both levels of information, using experimental observables to refine and validate the simulated ensembles [39, 40, 45–47]. In the case of smFRET, distance restraints constrain the accessible conformational space and increase the precision of the resulting structures [47–49], an approach well established for proteins [50] and becoming increasingly relevant for RNA [35, 39, 51].

Translating structural ensembles and local folding trajectories into FRET observables requires an appropriate representation of the attached fluorophores including relative dye-dipole orientations and photophysical parameters such as donor’s quantum yield etc. [28, 52, 53]. This is particularly relevant for cyanine fluorophores, whose interactions with the local RNA environment can restrict dye mobility and alter their photophysical properties [54, 55]. Such environment-sensitive responses includes protein- or RNA-induced fluorescence enhancement, in which changes in dye isomerization, stacking, and steric restriction modulate dyes fluorescence emission [56] and should be considered for accurate FRET predictions [51, 57, 58].

In MD simulations, fluorescent dyes are included explicitly as all-atom representation [42, 55, 58–60] or approximated as post-hoc dye model by the sterically accessible volume (AV) [61, 62] together with the contact volume (CV) of dye and linker attached to the host-molecule [55]. The latter yielding multiple accessible contact volumes (mACVs) for fluorescently labeled structural ensembles or along MD trajectories, respectively [35, 39, 50]. Therein, the dynamic fluorescence anisotropy serves as experimental measure for the dye mobility by weighting the CV relative to the AV [39]. In this way, dye–RNA interactions improve the comparability of simulated/predicted and experimental FRET observables [50].

In previous work, we characterized the ribosomal RNA model construct of the TLR in its unbound state, in which the H22/H88 KL is formed in the presence of monovalent metal ions, while the TL_GAAA_ remains unbound [35]. For this purpose, representative structures were sampled in a FRET-guided selection approach from a predicted structure collection. Here, we now focus on the bound state, in which the H22/H88 KL and the H68 TL_GAAA_ engage in a defined tertiary arrangement in the presence of divalent metal ions. We applied FAMP [51], our semi-automated FRET-assisted integrative modeling pipeline, which combines RNA 3D prediction, *in silico* fluorophore labeling, MD simulations, and prediction of fluorescence observables within a single workflow. We assembled the bound state in a knowledge-based manner from high-resolution ribosomal RNA structural elements and probed its stability by restrained and unrestrained all-atom MD simulations. These simulations sampled local rearrangements of the bound state and distinct fluorophore– RNA interactions. Dye stacking, transient contacts, and linker-dependent dynamics directly influence the apparent FRET efficiency and allow a molecular interpretation of the measured smFRET distributions. Together, the previously characterized unbound ensemble and the bound-state ensemble established here define the two limiting cases of the Mg^2+^-dependent folding equilibrium. By recording smFRET distributions across a range of Mg^2+^ concentrations and relating intermediate FRET states to different contributions of these reference ensembles, we reconstruct the structural transition from the unbound to the bound KL– TL_GAAA_ contact.

## Methods

The FRET-assisted modeling pipeline (FAMP) integrates multiple software components for integrative structure modeling of nucleic acids described recently [51]. Therein, four Python-based software modules — `Modeling, MD simulation, FRET measures` and `Evaluation` — cover the full path of FRET-guided integrative modeling. Initialized with a nucleic acid sequence FAMP produces FRET and dynamic fluorescent anisotropy predictors based on fluorescently labeled 3D RNA structures summarized in Figure 1 and SI Figure 1. The semi-automated pipeline can be run iteratively, and each step remains accessible to user intervention, allowing different modeling and simulation strategies to be assessed rapidly.

**Figure 1.**
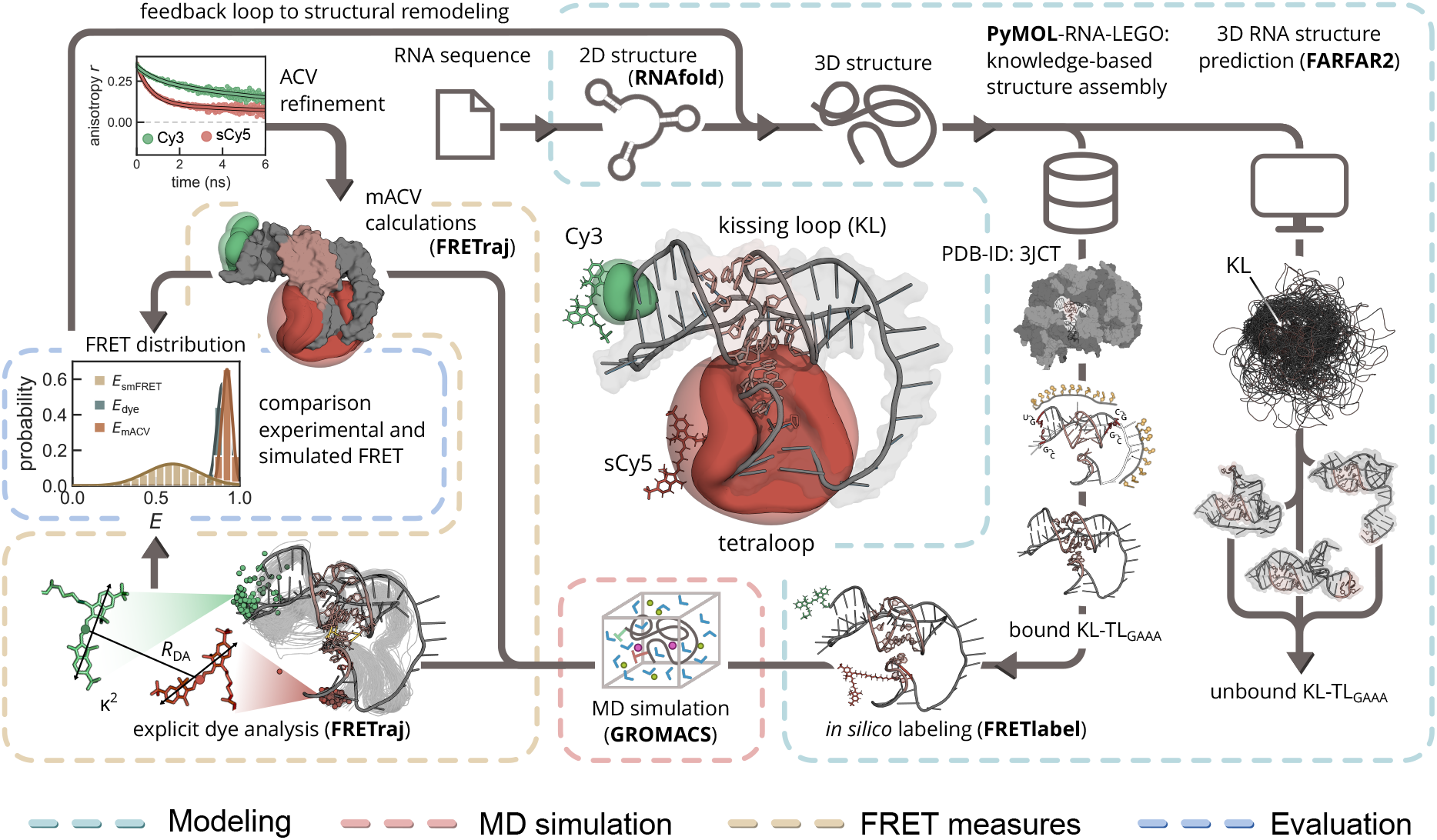
Workflow to model the KL–TL_GAAA_ tertiary contact in FAMP. Dashed frames indicate the four FAMP modules: Modeling (light blue), MD simulation (red), FRET measures (yellow), and Evaluation (dark blue). Starting from the RNA sequence and its predicted secondary structure, two routes led to a three-dimensional RNA model: knowledge-based assembly using structural elements of the cryo-EM reference structure (PDB ID: 3JCT) as a starting point, giving the bound KL–TL_GAAA_ state (bottom center), and *de novo* RNA 3D structure prediction using Rosetta’s FARFAR2 (right). Both models are subsequently labeled *in silico* with Cy3 and sCy5 using FRETlabel and passed to all-atom MD simulations using GROMACS. FRET observables are predicted from the resulting MD trajectories along two parallel branches: using explicit all-atom dyes, from which the inter-dye distance *R*_DA_ and the orientation factor *κ*^2^ are extracted (bottom left), and the mACV dye model (center left), where the contact-volume fraction is defined by experimental anisotropy decays of each dye (top left). Eventually, the predicted FRET distributions *E*_dye_ and *E*_mACV_ are compared with the experimental FRET histogram *E*_smFRET_.

### Experimental fluorescence observables

FRET measurements were performed on the Cy3/sCy5-labeled ribosomal RNA model construct purchased from Ella Biotech, in order to monitor TL_GAAA_ binding to the H22/H88 KL [14]. All measurements were carried out at room temperature in standard buffer (20 mM HEPES, pH 7.5, 100 mM KCl) at MgCl_2_ concentrations ranging from 0 mM to 100 mM. Prior to measurement, RNA samples were heated to 70 ^*°*^C for 2 min to unfold the RNA. Ensemble FRET measurements were performed at an RNA concentration of 100 nM using a SpectraMax iD5 multimode microplate reader (Molecular Devices). Recorded fluorescence intensities were corrected for background fluorescence and spectral cross-talk before FRET efficiencies were calculated (SI Methods).

Single-molecule FRET measurements were performed using a MicroTime 200 confocal fluorescence microscope (PicoQuant) operated in pulsed interleaved excitation (PIE) mode in HybriWell microfluidic chambers (Grace Biolabs). To minimise non-specific surface binding, the chambers were incubated with 1 mg mL^*™*1^ bovine serum albumin (Carl Roth) for 10 min and subsequently rinsed with standard buffer. RNA samples were loaded at a final concentration of approximately 100 pM. The donor and acceptor fluorescence were recorded in separate spectral detection channels for two hours each. Single-molecule fluorescence data were corrected for background fluorescence, spectral cross-talk, and differences in detection efficiencies and condition dependent fluorophore quantum yields (SI Table S1 and S2), *i*.*e*., *γ*-correction [53], and further analyzed using PAM [63]. Bursts were identified using an all-photon burst search. ALEX-2CDE filtering was additionally applied to exclude donor-only and acceptor-only species and to reduce the contribution of bursts affected by fluorophore blinking or photobleaching [64] (SI Methods and SI Figure S2).

Fluorescence dynamic anisotropy decays were recorded using a FluoTime 250 fluorescence lifetime spectrophotometer (PicoQuant) equipped with a Prima-450-515-640 ps laser and a PMA Hybrid 07 single-photon counting detector. RNA samples in the respective standard buffer containing varying MgCl_2_ concentrations were excited at 511 nm and 640 nm with a repetition rate of 15 MHz, and the polarization-resolved fluorescence emission was collected using 570/60 nm and 700/70 nm bandpass filters, respectively.

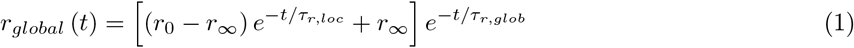

with *r*_0_ describing the fundamental anisotropy, *r*_*∞*_ the residual anisotropy, *τ*_*r,loc*_ as the local rotation time of the dye in the realm of the nucleic acid, and *τ*_*r,glob*_ as the global rotation time of the entire construct. The dye-nucleic acid stacking probabilities (*χ*) were calculated according to [39] (SI Methods, SI Figure S3 and S4, SI Table S5)

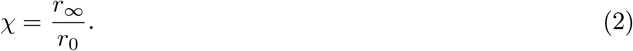

### 3D RNA structure modeling

Using the Modeling module of FAMP, we generated the bound conformation of the KL-TL model construct via knowledge-based structure assembly. Therein, structural elements from experimentally resolved 3D RNA structures were used. The model construct described by Gerhardy et al. [14] provided the target RNA sequence and construct architecture, while the H22/H88 TLR with the bound H68 TL_GAAA_ was extracted from the cryo-EM structure reported by Wu et al. [18] (PDB-ID: 3JCT) to preserve the KL–TL_GAAA_ tertiary-contact geometry (Figure 2A). Individual bases in the H22 and H88 stems were then mutated in PyMOL to match the Gerhardy et al. model construct sequence. The three helices were subsequently connected by inserting poly-A linker elements using the PyMOL Builder functionality. As a result, we obtained an RNA structural model of the bound construct that preserves the cryo-EM-derived KL–TL_GAAA_ contact while matching the sequence and linker architecture of the KL–TL_GAAA_ experimental construct (Figure 2B). Subsequent *in silico* labeling of the RNA construct at the experimentally defined positions (Figure 2C). The internal sCy5 label was introduced using FRETlabel [39], retaining the AMBERDYES-compatible nomenclature required for subsequent MD simulations [59]. The specific Cy3 dye–linker configuration at the 3’-end was attached manually in PyMOL (SI Methods).

**Figure 2.**
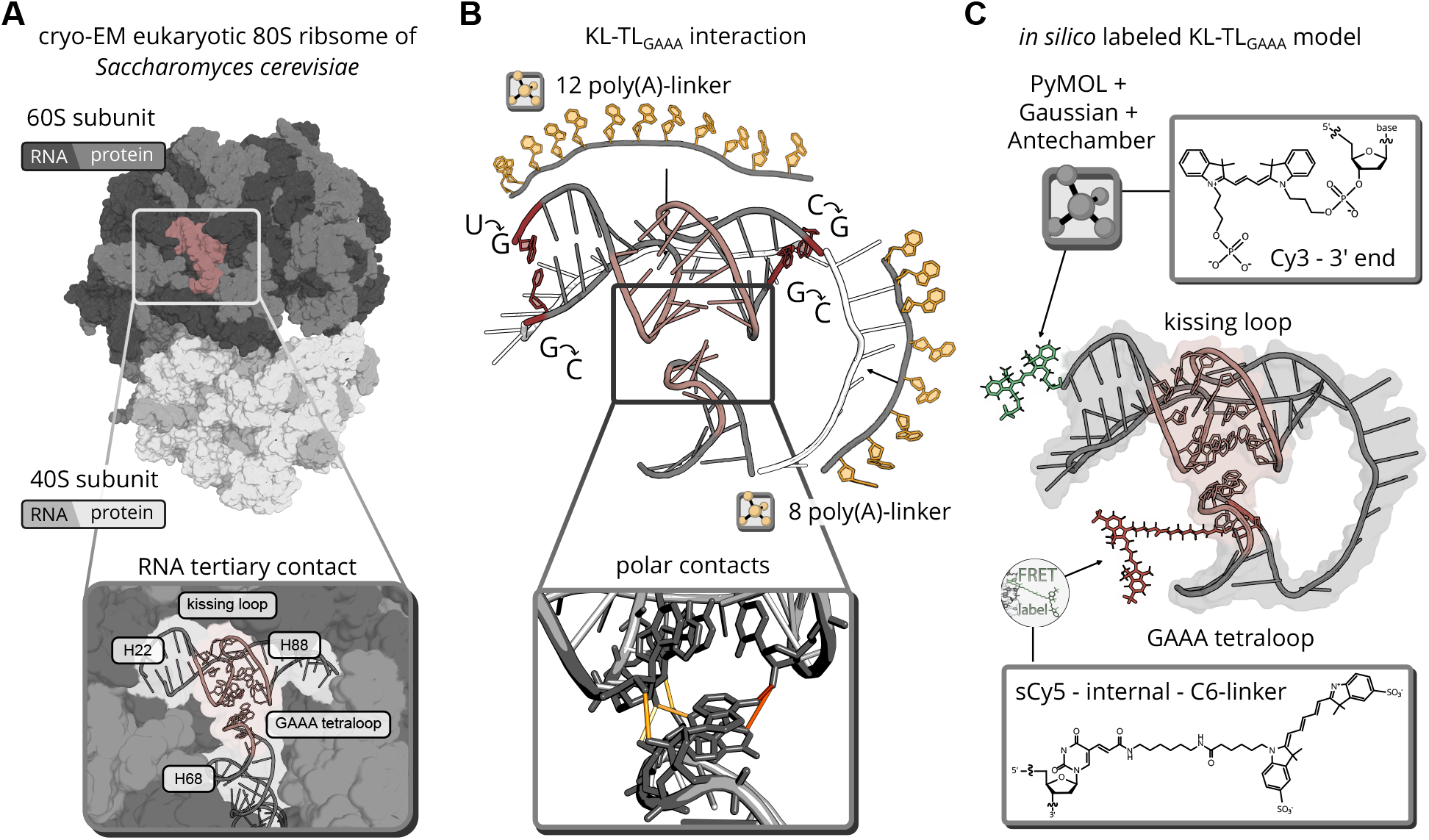
Modeling workflow from the cryo-EM structure to a dye-labeled RNA 3D model preserving the experimentally resolved bound conformation. **(A)** Cryo-EM structure of the yeast large ribosomal subunit (PDB ID: 3JCT), showing the long-range tertiary contact between the H68 TL_GAAA_ and the H22/H88 KL, which acts as the TLR. **(B)** Structural elements extracted from the ribosomal subunit are shown in grey. Poly-A linkers modeled in PyMOL are shown in yellow, while the cryo-EM-derived non-covalent interactions between the TL_GAAA_ and the TLR are preserved. **(C)** *in silico* attachment of Cy3 and sCy5 to the RNA model construct. Cy3 was attached to the 3^*′*^ end via PyMOL, whereas sCy5 was labeled internally with FRETlabel to monitor conformational changes associated with TL_GAAA_ binding. The corresponding dye structures are shown in gray boxes.

### MD simulations

All-atom MD simulations of the *in silico* rRNA model construct were performed using GROMACS release 2025.3 [65]. The RNA was described by the AMBER force field [66] with bsc0 [67] and *χOL*3 corrections [68, 69]. Parameters for sCy5 were taken from AMBER-DYES [59, 70]. For Cy3, the fluorophore and linker chemistry corresponding to the commercially synthesized RNA construct were parameterized specifically for this study according to established procedures for MD simulations (SI Methods). This parameterization provides a transferable basis for extending existing dye parameter sets and labeling libraries to fluorophores and linker chemistries that are now routinely available from commercial suppliers. A dye fragment reproducing the actual attachment chemistry was built *in silico* using PyMol [71] and parametrized by geometry optimization and ESP calculation in Gaussian followed by a RESP fit with AmberTools [72]. Structures were solvated in explicit TIP4P-Ew water [73]. K^+^ ions were added to neutralize the negatively, together with Cl^-^, to a final KCl concentration of 100 mM. Mg^2+^ ions were placed randomly at final concentrations of 20 mM and 100 mM, with the corresponding ion and force-field parameters specified in the SI (SI Methods). Four production MD runs of 1 *µ*s each were performed, combining the two Mg^2+^ concentrations with the presence or absence of distance restraints in order to keep the initial cryo-EM structure of the ribosomal RNA KL-TL interaction intact (SI Table S4 and SI Table S5). Restraints were applied as pairwise linear/harmonic distance restraints derived from the interaction of the original cryo-EM construct (SI Methods and SI Figure S5).

### *in silico* FRET and dynamic anisotropy calculation

FRET observables were calculated from the MD trajectories using two complementary fluorophore representations, explicit all-atom dyes and the mACV model. Both representations were evaluated within the DataAnalysis module of the FAMP pipeline [51], using MDAnalysis [74, 75] and FRETraj [58] (SI Methods and SI Table S6). Time-dependent donor–acceptor distances *R*_DA_(*t*) and orientation factors *κ*^2^(*t*) were used as input for photon-burst simulations with FRETraj Burst (SI Methods). For the unbound state, we followed the previously established FRET-guided selection and burst-simulation scheme, in which individually selected conformations contribute according to their selection probabilities, assuming that RNA conformational changes occur on timescales longer than the duration of individual bursts in the smFRET experiment [35]. Donor and acceptor photon emission events, *I*_D_ and *I*_A_, were generated using a Markov chain Monte Carlo scheme to obtain simulated FRET distributions and dynamic fluorescence anisotropy decays directly comparable to the experimental fluorescence data [42, 58]. Therein, photon events were simulated based on the experimentally determined fluorescence lifetimes of *τ*_*D*_ = 1.67 ns for Cy3 and *τ*_*A*_ = 1.42 ns for sCy5. For each experimental condition, 20,000 photon bursts were simulated using the corresponding experimental burst-size distribution together with the [Mg^2+^]-dependent fluorescence lifetimes *τ*_D*/*A_, detection efficiencies *η*_D*/*A_, and fluorescence quantum yields *Q*_D*/*A_ of the donor and acceptor fluorophores (SI Figure S6). The resulting *in silico* FRET efficiencies were *γ*-corrected using the condition-specific experimental parameters to enable direct comparison with the fully corrected experimental FRET distributions (SI Methods and SI Table S2 and S7-S8).

### Population-weighted mixing of dye representations and unbound/bound ensembles

Population-weighted mixing was used both to combine the explicit-dye and mACV representations of the bound state and, subsequently, to combine the unbound and bound conformational ensembles. In both cases, mixtures were generated directly during photon-burst simulation using `averaging = “ensemble”`.

Each input ensemble was treated as a separate species (*S*) with an assigned population weight *w*_*i*_, such that

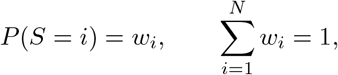

where *P* (*S* = *i*) denotes the probability of selecting species *i*. In FRETraj’s `ensemble`-averaging mode, the species was resampled according to these probabilities for every relaxation event within a simulated burst. Thus, a mixture with weights *w*_A_ = 0.85 and *w*_B_ = 0.15 corresponds to repeated sampling from the complete ensembles A and B with probabilities of 85 % and 15 %, respectively.

Within each selected species, the corresponding trajectory was used directly for the respective relaxation event. As each species was represented by a single trajectory, no additional trajectory-level weighting was required. Population weights were analyzed systematically from 0 to 1 in increments of 0.05, *w ∈* {0, 0.05, 0.10, *…*,1.00}. The weighting that yielded a simulated mean FRET efficiency closest to the corresponding experimental smFRET mean for the particular Mg^2+^ concentration chosen was selected as fixed dye representation of the particular conformational state or its ensemble.

## Results

### Cryo-EM based MD simulations reflect FRET measurements at supraphysiological Mg^2+^ concentrations

The structural baseline for our modeling pipeline comprised the TLR and TL fragments derived from the cryo-EM structure [18] and adapted in sequence and linker architecture to match the FRET construct described by Gerhardy et al. [14]. First, we calculate donor-acceptor distances from this knowledged-based 3D RNA structure model of the KL–TL_GAAA_ in its bound state. ACV calculations yielded an average donor–acceptor distance of 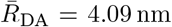 nm, corresponding to an energy transfer efficiency of *Ē*_ACV_ = 0.89 (Figure 3A). This ACV-based first estimate is significantly higher than the observed fully corrected mean FRET value *Ē*_smFRET_ = 0.79 ± 0.12 under experimental conditions of 116 mM K^+^ and an unphysiologically high Mg^2+^ concentration of 100 mM (Figure 3B).

**Figure 3.**
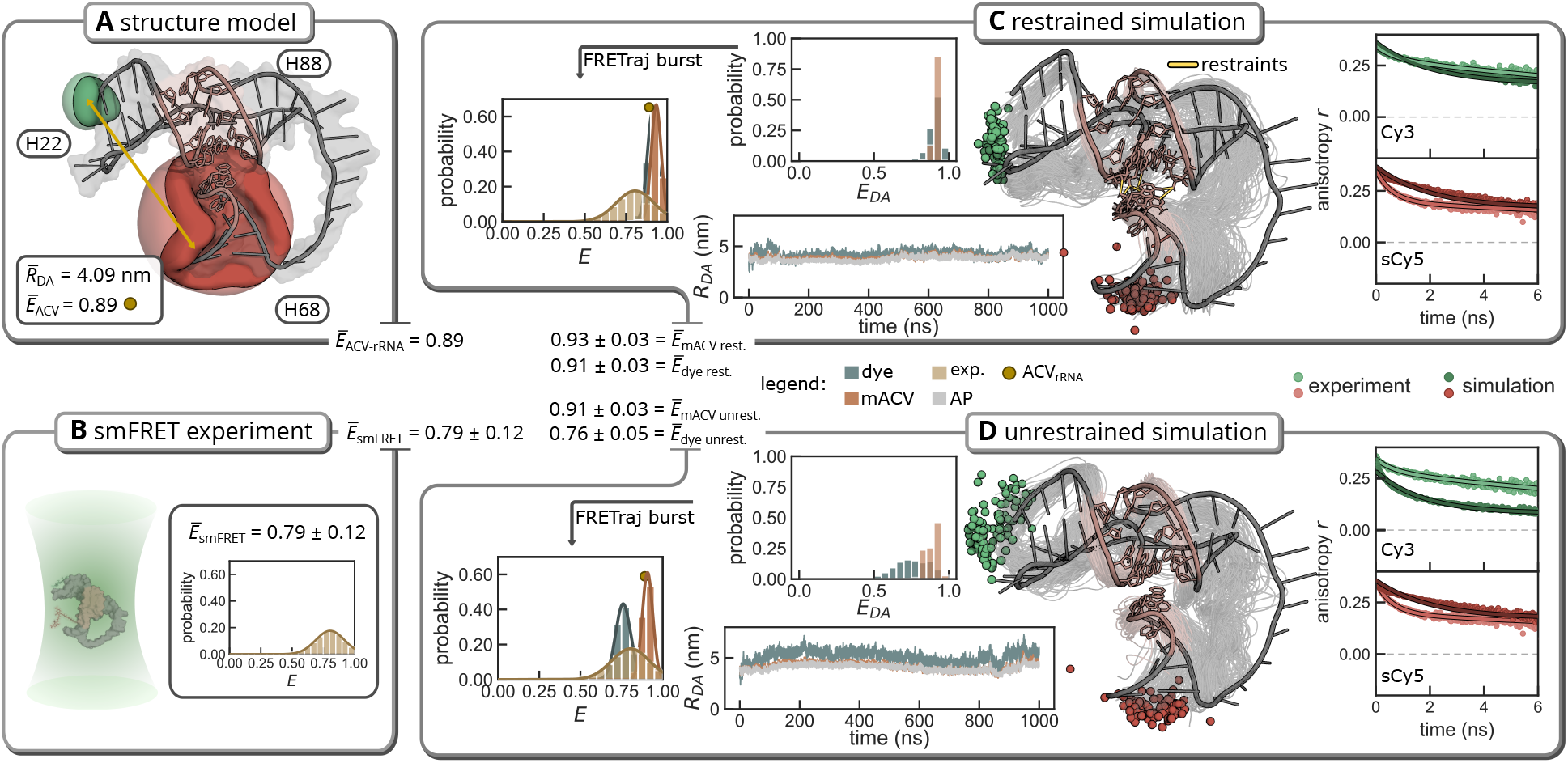
Comparison of experimental smFRET data with a single structure estimate and restrained and unrestrained MD structure collections. **(A)** Modeled bound state of KL–TL_GAAA_ showing the predicted ACVs, the mean donor–acceptor distance R_DA_, and the corresponding E_ACV_ value. **(B)** Experimental smFRET distribution at an Mg^2+^ concentration of 100 mM, including the corresponding mean and standard deviation. Both **(C)** restrained MD simulation and **(D)** unrestrained MD simulation show the mean C-atoms of the respective dyes Cy3 and sCy5, together with the dye-, attachment-point-, and mACV-derived R_DA_ distances shown in the lower left. The distance-derived E_DA_ distribution is shown in the upper left, whereas the photon-burst-derived FRET histogram, which enables direct comparison with the experiment, is shown on the left for each simulation. Experimental anisotropy measurements and comparison to the all-atom dye simulations are shown on the right for both simulations.

To simulate more realistic smFRET experiments we resampled structural and dye conformations from the bound RNA model construct. To this end, we performed 1 µs MD simulations starting from the cryoEM-derived KL–TL_GAAA_ model, which allowed us to assess both stability and dynamics of the bound RNA tertiary contact and the mobility of the attached fluorophores in the RNA environment. We compared two complementary simulation strategies. In the restrained simulation (Figure 3C - RNA model), distance restraints were applied to maintain the KL–TL_GAAA_ contact in its cryo-EM-derived bound geometry (SI Figure S5C), thereby mimicking a confined native-like state as stabilized within the ribosomal context. In contrast, the unrestrained simulation allowed the isolated TL to relax freely and possibly undock from the TLR as in the *in vitro* solution experiment (Figure 3D - RNA model). Both 1 µs restrained and unrestrained simulations yielded highly similar structural ensembles in which the KL–TL_GAAA_ interaction remained intact. The donor–acceptor distances *R*_DA_ derived from the mACVs and dye attachment points (AP) were stable throughout both MD trajectories. In the restrained simulation, the mean mACV-derived distance was *R*_DA_ = 4.22 nm ± 0.35 nm, yielding a mean values of *Ē*_DA_ = 0.91 ± 0.04. In the unrestrained simulation, mean mACV-derived distance of *R*_DA_ = 5.11 nm ± 0.58 nm produced a FRET efficiency of *Ē*_DA_ = 0.89 ± 0.04.

Next, photon-burst simulations were performed for each trajectory based on the mACV-dye model and explicit all-atom dyes. This approach not only accounts for the shot noise broadening of the FRET distribution using the particular burst size distribution of the experiment, but also includes gamma factor and cross-talk corrections (SI Methods, SI Figure S6), enabling a direct comparison between predicted and experimentally observed transfer efficiency distributions (SI Table S8). For the restrained simulation, the all-atom dye trajectory yielded a mean FRET efficiency of *E*_dye rest._ = 0.91 ± 0.03; the corresponding mACV-based calculation yielded *E*_mACV rest._ = 0.93 ± 0.03. Here, both predicted FRET distributions fully overlap with each other, but differ in mean and width from the experimental FRET distribution. For the unrestrained simulation, the mACV-derived distances similarly produced a high FRET efficiency of *E*_mACV unrest._ = 0.91 ± 0.03, equal to that of the restrained simulation. However, the larger donor–acceptor distances sampled by the explicit all-atom dye model shifted the corresponding distribution towards lower FRET efficiencies, resulting in *E*_dye unrest._ = 0.76 ± 0.05. Thus, the unrestrained all-atom dye trajectory not only yielded a mean FRET efficiency close to the experimental smFRET value, but both models together cover the experimental FRET distribution.

While the mACV-based FRET predictions remained essentially unchanged at *E*_mACV_ *>* 0.9, the explicit-dye trajectories sampled two distinct fluorophore configurations despite the overall RNA structure remaining similar in the restrained and unrestrained simulations. In the restrained trajectory, Cy3 remained positioned closer to the sCy5 acceptor, consistent with the high-FRET regime also predicted by the mACV representation. By contrast, the unrestrained trajectory sampled an alternative Cy3 interaction at a larger donor–acceptor separation, shifting the explicit-dye FRET distribution towards lower FRET efficiencies of *E*_dye unrest._ = 0.76 ± 0.05. Thus, the difference between the restrained and unrestrained FRET predictions arises predominantly from fluorophore behavior rather than from major rearrangements of the RNA scaffold. These two limiting dye configurations broaden the range of predicted FRET efficiencies and together span the experimentally observed FRET distribution.

### Dye stacking partitions the bound state into spectroscopically distinct subpopulations

To assess whether restrained and unrestrained MD simulations indeed recall experimentally measurable fluorophore dynamics on the host RNA molecule, dynamic fluorescence anisotropy decays were calculated from all-atom dye MD trajectories and compared to the experiment at the particular [Mg^2+^] = 10 mM and 100 mM (Figure 3C and D (100 mM) and SI Figure 7C and D (10 mM)).

Interestingly, the dynamic anisotropy decay comprises two separate decay components (SI Table S7). The fast component *τ*_*r,loc*_ is attributed to the dye movement within a cone given by the dyes geometrical properties and the actual sterical restriction of the host molecule. Here, Cy3 and sCy5 show experimental values *τ*_*r,loc*_ *<* 1*ns* which are in the upper ps range and in general comparable with the simulated decays. However, all simulated decay times *τ*_*r,loc*_ were faster than in the experiments, which is indicative for a stronger restriction of the dye movement *in silico* than actually reported in the experiment. Along this line, the experimental residual *r*_*∞*_ and fundamental *r*_0_ anisotropy for Cy3 calculated to a stacking probability of *χ* =82.5 %. Here, the restrained simulation yielded a similar high stacking probability of 92.5 %, whereas the unrestrained simulation yielded a lower value of 54.7 %. Consistent with these differences, fitting the wobbling-in-a-cone model gives cone semiangles *θ*_0_ = 20.3° for the experiment, *θ*_0_ = 12.9° for the restrained simulation, and *θ*_0_ = 35.4° for the unrestrained simulation. Thus, under the investigated [Mg^2+^] condition, the restrained trajectory reproduced the experimentally observed dye mobility restriction of Cy3 more closely than the unrestrained trajectory.

For sCy5, both simulations showed substantially stronger stacking than observed experimentally. The experimental residual *r*_*∞*_ and fundamental *r*_0_ anisotropy yielded a stacking probability of 56.4 %, whereas the restrained and unrestrained simulations resulted in stacking probabilities of 93.5 % and 90.4 %, respectively. The corresponding cone semiangles further emphasized this discrepancy, with *θ*_0_ = 34.6° obtained experimentally compared with *θ*_0_ = 12.0° and *θ*_0_ = 14.8° for the restrained and unrestrained simulations, respectively. The structural 3D RNA structure ensembles with explicit all-atom dyes showed that the restricted rotational freedom resulted from persistent stacking interactions of sCy5 at the 5^*′*^ end of H68. Although the unrestrained simulation permitted slightly greater sCy5 mobility than the restrained simulation, neither trajectory reproduced the conformational freedom observed experimentally.

The pronounced difference in stacking probabilities of both dyes therefore indicate that fluorophore dynamics are strongly influenced by the local RNA environment, i.e., the particular labeling site. Despite the lower stacking probability of Cy3 observed in the unrestrained simulation, the reduced stacking of the dye resulted in FRET efficiencies that more closely matched the experimental smFRET data, as reflected in both the distance-derived *E*_DA_ values and the photon-burst FRET distributions. In contrast, sCy5 was substantially more restricted in both simulations than observed experimentally, despite the C6 linker providing greater conformational freedom than the linker-free Cy3 attachment. The pronounced sCy5 stacking in the simulations therefore presents a clear deviation from the experimentally observed dye dynamics. Together with the comparatively short Cy3 linker, pronounced dye–RNA interactions likely contribute to the highly stable *R*_DA_ trajectories and the correspondingly narrow simulated *E*_DA_ distribution. Collectively, we posit that the experimentally measured FRET distribution is broadened by fluorophore dynamics, with both donor and acceptor dyes transiently stacking onto the RNA.

### Mg^2+^-dependent GAAA tetraloop binding is described by mixtures of *in silico* bound and unbound conformations

To characterize the Mg^2+^-dependent folding of the KL–TL_GAAA_ construct, we next performed FRET measurements at varying Mg^2+^ concentrations ranging from 0 to 100 mM (Figure 4 top). Higher Mg^2+^ led to an increase in FRET efficiency with an apparent association constant of *K*_*A*_ = 8.4 mM in an ensemble type experiment as reported before [14]. To structurally interpret this Mg^2+^-dependent two-state transition, we first used the conformational ensembles at the two limiting cases of the titration, i.e. bound and unbound, as reference states. We performed single-molecule FRET measurements and compared the fully corrected and filtered experimental FRET distributions with *in silico* FRET distributions based on RNA 3D structure collection for the particular condition (Figure 4, bottom).

**Figure 4.**
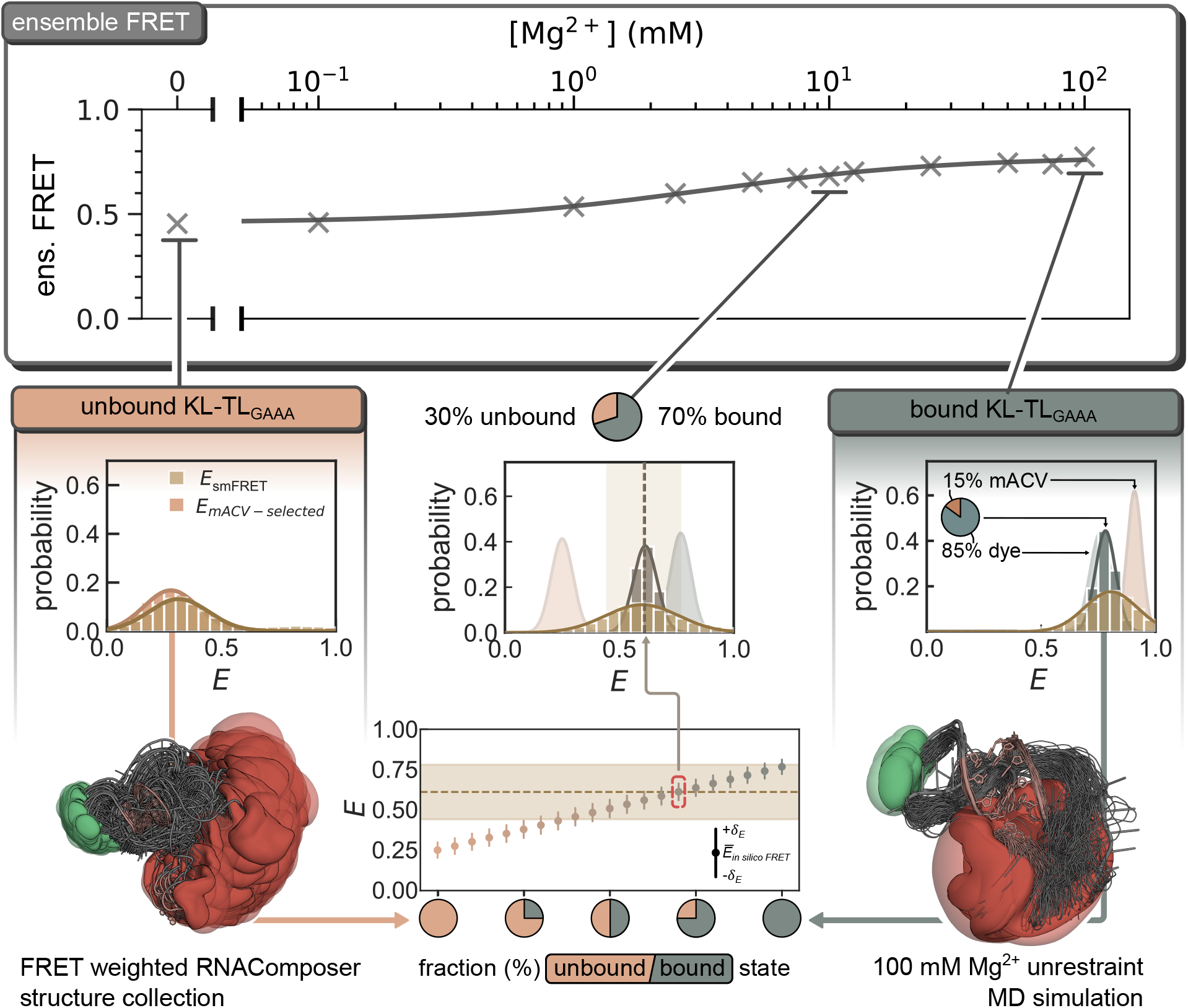
Mg^2+^-dependent ensemble and single-molecule FRET compared with *in silico* models of the bound and unbound states. **(Top)** Ensemble FRET measurements of the KL–TL_GAAA_ construct at Mg^2+^ concentrations ranging from 0 mM to 100 mM. **(Bottom)** smFRET distributions measured at [Mg^2+^] = 0 mM (left), 10 mM (middle), and 100 mM (right). The unbound-state smFRET distribution and corresponding structural ensemble were adapted from Weber et al., whereas the bound-state distribution and structural ensemble were determined in this study. To represent the intermediate smFRET distribution observed at [Mg^2+^] = 10 mM, the unbound and bound states were combined in a two-state model using their respective fractional populations.

Without divalent metal ions [Mg^2+^] = 0 mM we found the mACV-based *in silico* FRET distribution perfectly overlapping with the experimentally measured smFRET distribution (Figure 4 left, unbound KLTL_GAAA_). Herein, FRET-prediction was performed from a FRET-guided selection of an underlying RNA-Composer structure collection without averaging. Thus, a large variety of structures was selected to describe the *in silico* FRET distribution, whereas each individual structure contributed with their experimentally determined weight from the FRET probability density. This approach considers the transition between those structures, to be slow compared to the measurement time, *i. e*. the burst duration of the confocal fluorescence microscope.

With divalent metal ions at supraphysiological [Mg^2+^] = 100 mM we found both the mACV- and all-atom dye-based *in silico* FRET distributions overlapping the experimental FRET distribution but well separated with the experimental mean FRET centered between them. This good agreement indicates that the modeled bound conformation indeed represents unphysiological high-Mg^2+^ conditions or a confined local environment within the ribosome. The shift between both high-FRET distribution occurs, as the mACV approach does less weigh the CV than one would expect from the the all-atom dye MD trajectories. This is due to the experimentally observed more isotropic rotation of Cy5 and a lower stacking component compared to Cy3. However, we interpret the difference as limiting cases for Cy3/5 dye-stacking behavior on the RNA host molecule, *i. e*. stacked vs. (more) isotropic rotating, and we sampled mixtures of mACV- and explicit dye-derived distances to render the actual experimental results. Therein, a combination of 85 % explicit dye distances and 15 % mACV-derived distances were averaged and reproduced the mean of the experimental smFRET distribution best (Figure 4, right, bound KL-TL_GAAA_). However, the width of the distribution could not be reproduced (SI Figure S8). This agreement suggests that Cy3 may populate different interaction states within its accessible RNA environment, including both more weakly associated and stacked configurations, thereby contributing to the broader experimental smFRET distribution.

From the ensemble FRET experiment, we are aware of the transition between two states, the bound and unbound conformation of the KL-TL construct. Comprising a *K*_*A*_ = 8.4 mM. We choose a close value of [Mg^2+^] = 10 mM to probe the transition between both states on the single molecule level and found a broad FRET distribution between the two limiting cases described above. Indeed, the intermediate FRET efficiency suggested that both unbound and bound conformations contributed to the experimental ensemble. As we found structurally defined references for the low- and high [Mg^2+^], these two experimentally validated structure collections can be regarded as references across the Mg^2+^ titration. Using these structurally defined states, we systematically varied the relative fractions of bound and unbound conformations to account for the conformational heterogeneity and calculated the corresponding photon-burst FRET distributions (Figure 4, middle). Increasing the contribution of the bound-state ensemble progressively shifted the predicted FRET efficiencies toward the high-FRET regime. The experimentally observed Mg^2+^-dependent transition could therefore be described by changing relative populations of the two conformational ensembles, providing a structural interpretation of the folding behavior observed in the ensemble FRET titration. Notably, the smFRET distribution measured at [Mg^2+^] = 10 mM was best reproduced by a mixture containing approximately 70 % unbound and 30 % bound conformations (Figure 4, bottom middle) although with a smaller width of the distribution but perfectly matching the mean FRET value observed experimentally.

## Discussion

FRET-guided integrative modeling, particularly when combined with smFRET experiments, provides a powerful framework for moving beyond static structural representations toward ensembles of interconverting or coexisting conformations [48, 50, 76]. This ensemble perspective has become increasingly important in RNA structural biology, where function often depends on the relative populations of multiple conformational states rather than on a single lowest-energy structure [24, 39, 77]. Extending a single structural representation toward such conformational ensembles through simulations is possible but, involves several methodological choices and potential pitfalls that can strongly influence the resulting structural interpretation [35, 40]. Although smFRET provides direct access to conformational (sub-)populations, similar FRET efficiencies can arise from different RNA conformations, fluorophore positions, or dye–RNA interactions, making their structural assignment inherently ambiguous [13, 78, 79]. Resolution of such degenerate FRET states requires careful inspection of orthogonal measures, such as anisotropy, spatial restraints or alternative labeling pairs. In the present study, these methodological considerations proved critical for resolving the structural relationship between the unbound and bound KL–TL_GAAA_ ensembles.

For the KL–TL_GAAA_ system, we therefore describe the folding landscape using experimentally informed reference ensembles, as a comprehensive atomistic description of the underlying conformational landscape remains challenging with currently accessible sampling approaches [43, 80]. Importantly, the two structural limits are not equally accessible to current RNA 3D prediction methods. The flexible unbound ensemble could be resolved by FRET-guided selection from predicted structure collections, as previously demonstrated by Weber et al.[35], whereas the long-range KL–TL_GAAA_ contact remained inaccessible to all tested prediction approaches. This disparity reflects a broader limitation of RNA structure prediction, where long-range non-Watson–Crick tertiary interactions and pseudoknot-like topologies remain substantially more difficult to capture than canonical secondary-structure elements, as repeatedly observed in RNA-Puzzles [81, 82]. The cryo-EM-derived bound model should therefore be regarded as an experimentally informed structural hypothesis that provides a defined starting point for MD simulations to explore the conformational space, stability, and structural variability of the construct.

Using this cryo-EM-derived model as a common structural starting point, the restrained and unrestrained MD simulations provide complementary views of the conformational space accessible to the bound KL– TL_GAAA_ state. Both simulations suggest that the bound KL–TL_GAAA_ state should not be viewed as a single well-defined conformation, but rather as an ensemble whose FRET efficiency depends on both local RNA structure and fluorophore dynamics. Preservation of the cryo-EM-derived geometry consistently favored a compact high-FRET state, largely independent of the fluorophore representation, indicating that this geometry likely corresponds to a highly compact subpopulation of the bound ensemble in solution. Allowing the construct to relax without structural restraints broadened the accessible conformational space while preserving the tertiary contact, demonstrating that noticeable changes in FRET can occur without dissociation of the KL–TL_GAAA_ interaction.

The conformational variability of the bound RNA alone, however, does not fully account for the differences between the predicted and experimental FRET distributions. A substantial contribution arises from the fluorophores and their interactions with the local RNA environment. The difference between mACV- and explicit-dye predictions illustrates this directly, with explicit dyes sampling lower FRET efficiencies in the unrestrained trajectory while the mACV representation remained shifted toward higher values. The experimental mean could be reproduced by combining 85 % explicit dye- and 15 % mACV-derived distances, consistent with heterogeneous fluorophore configurations contributing within the relaxed bound-state ensemble. The anisotropy analysis provides an independent constraint on this interpretation. While the simulated Cy3 dynamics approaches the experimentally observed degree of restriction, sCy5 remains substantially more stacked and rotationally restricted than observed experimentally, despite its comparatively flexible linker. Agreement between calculated and experimental FRET efficiencies alone is therefore insufficient to validate a particular structural ensemble. Instead, the combined FRET and anisotropy data indicate that local dye–RNA interactions form an integral part of the measured observable and can substantially modulate the apparent FRET efficiency. The remaining discrepancies, particularly for sCy5, likely reflect limitations in the representation and sampling of fluorophore–RNA interactions in addition to conformational heterogeneity of the RNA itself [28, 56].

Despite this sensitivity of the FRET observable to local fluorophore dynamics, the structurally characterized unbound and bound ensembles provide a useful framework for interpreting the Mg^2+^-dependent folding landscape. The broad smFRET distribution of the previously characterized unbound ensemble at low Mg^2+^ indicates that its conformational heterogeneity is not fully averaged within the observation time of an individual burst, consistent with an exchange between distinct unbound conformations that is similar to the burst duration. At the opposite end of the titration, the bound state is likewise better described as an ensemble of locally distinct conformations and fluorophore configurations than by a single cryo-EM-like structure. The Mg^2+^-dependent transition is therefore described here as a redistribution between unbound and bound populations, while both states retain substantial internal conformational heterogeneity.

The interpretation of intermediate Mg^2+^ conditions is further complicated by the fact that several contributions to the measured FRET efficiency change simultaneously and are thermodynamically coupled. Increasing Mg^2+^ promotes electrostatic screening and RNA compaction [16, 83, 84], favors formation of the KL–TL_GAAA_ tertiary contact [12, 85], and can additionally modify the local environments and interactions of the attached fluorophores, thereby affecting their photophysical properties, including the quantum yield [28, 55]. Changes across the FRET titration therefore cannot be assigned exclusively to formation of the tertiary contact itself. Nevertheless, weighting these reference ensembles provides a simple structural description of the progressive Mg^2+^-dependent shift. At [Mg^2+^] = 10 mM, the experimental smFRET distribution is best represented by approximately 70 % bound and 30 % unbound conformations, despite KL–TL_GAAA_ binding having previously been reported to approach saturation under these conditions. This distinction suggests that near-saturating binding does not imply predominant occupation of the structurally defined high-FRET bound ensemble. Instead, a substantial fraction of molecules can remain within unbound-like or near-bound regions of conformational space even when tertiary-contact formation is strongly favored.

Overall, the continued Mg^2+^-dependent evolution of the FRET signal beyond near-saturating binding indicates that formation of the KL–TL_GAAA_ contact does not define a unique structural endpoint, but remains coupled to RNA compaction, conformational heterogeneity, and fluorophore response.

## Conclusion

In this study, FRET-guided integrative modeling was used to connect Mg^2+^-dependent smFRET states with structurally interpretable ensembles of the KL–TL_GAAA_ construct. Although the bound tertiary contact could not be resolved *de novo* by current RNA 3D prediction approaches, FAMP enabled the integration of knowledge-based structural assembly, restrained and unrestrained MD simulations, complementary fluorophore representations, photon-burst simulations, and fluorescence anisotropy within a common modeling framework. This combination allowed experimentally anchored unbound and bound reference ensembles to be established and related to the conformational heterogeneity observed in solution.

The Mg^2+^-dependent transition is therefore best described not as a conversion between two discrete structures, but as a redistribution between structurally heterogeneous unbound and bound ensembles. Intermediate FRET states can be represented by changing contributions of these reference ensembles, while local RNA rearrangements and fluorophore dynamics remain important determinants of the measured signal. In particular, the bound KL–TL_GAAA_ state comprises a broader conformational ensemble than the compact cryo-EM-like geometry alone, emphasizing that structural interpretation of smFRET data requires explicit consideration of both RNA conformational variability and dye–RNA interactions.

The structural framework established here provides a basis for the future integration of dynamic fluorescence observables such as smFRET to resolve the timescales and pathways of conformational exchange within and between these ensembles. Together, the experimentally anchored structural states and their Mg^2+^-dependent redistribution establish a foundation for extending FRET-guided integrative modeling toward a dynamic description of of the KL–TL_GAAA_ tertiary-contact formation.

## Competing interests

No competing interest is declared.

## Data Availability

The source code of FRETraj is available on GitHub https://github.com/RNA-FRETools/fretraj and Zenodo https://doi.org/10.5281/zenodo.10898652. Documentation can be found at https://rna-fretools.github.io/software/. Additional source material is available from the corresponding author upon request.

## Acknowledgements

F.E., M.W., V.S., M.H., J.M., and R.B. gratefully acknowledge financial support by the European Social Fund Plus for Germany (ESF, Grant No. 100649226 to R.B., PhD scholarship to M.H.), the German Research Foundation (DFG, Grant No. 498128362 and 537331625 to R.B.), the Mittweida University of Applied Sciences (MUAS), and the Laserinstitut Hochschule Mittweida for any further financial support. Further, J.M. gratefully acknowledges her PhD scholarships to support women in academia and research at MUAS within the Federal and State ‘Women Professors Programme 2030’, budget for gender equality measures at MUAS. This project is cofinanced from tax revenues based on the budget adopted by the Saxon State Parliament and cofunded by the European Union.

## Supporting information

Additional supporting data is provided in the supplementary.pdf

## Author Contribution

F.E. (Conceptualization [equal], Data curation [lead], Formal analysis [lead], Investigation [lead], Methodology [equal], Project administration [supporting], Software [lead], Validation [equal], Visualization [lead], Writing – original draft [equal], Writing – review and editing [equal]), M.W. (Conceptualization [equal], Data curation [supporting], Formal analysis [supporting], Methodology [equal], Project administration [supporting], Software [supporting], Validation [equal], Visualization [supporting], Writing – original draft [equal], Writing – review and editing [equal]), V.S. (Formal analysis [supporting], Investigation [supporting], Methodology [supporting], Validation [supporting], Writing – original draft [supporting], Writing – review and editing [supporting]), M.H. (Formal analysis [supporting], Investigation [supporting], Methodology [supporting], Validation [supporting], Writing – original draft [supporting], Writing – review and editing [supporting]), J.M. (Formal analysis [supporting], Investigation [supporting], Methodology [supporting], Validation [supporting], Writing – original draft [supporting], Writing – review and editing [supporting]), F.D.S. (Methodology [supporting], Software [supporting], Supervision [supporting], Writing – review and editing [equal]), P.K.Q. (Methodology [supporting], QM calculations [lead], Supervision [supporting], Writing – review and editing [equal]), R.B. (Conceptualization [lead], Formal analysis [supporting], Funding acquisition [lead], Investigation [supporting], Methodology [equal], Project administration [lead], Resources [lead], Supervision [lead], Visualization [supporting], Writing – original draft [equal], Writing – review and editing [equal]).

## Notes

### Competing Interest Statement

The authors have declared no competing interest.

