## Supplementary Information for "FRET-guided integrative modeling resolves Mg^2+^-dependent RNA tertiary-contact formation"

### Supplementary Methods

#### Determination of Correction Factors

##### Relative detection efficiency

The relative detection efficiency of the donor and acceptor detection channels of the MicroTime 200 was determined experimentally using a slope-based calibration adapted from McCann *et al.* (1), with the spectrofluorometer FluoroMax (Horiba) serving as the reference instrument. The relative detection efficiency accounts for differences in the wavelength-dependent transmission of the optical path and in the detector sensitivity of the donor and acceptor detection channels. It was calculated according to (1)

$$\eta_{A/D} = \frac{\eta_A}{\eta_D} = \left(\frac{m^D_{\text{FluoroMax}}}{m^D_{\text{MT200}}}\right)\left(\frac{m^A_{\text{MT200}}}{m^A_{\text{FluoroMax}}}\right), \tag{1}$$

where $\eta_D$ and $\eta_A$ denote the detection efficiencies of the donor and acceptor detection channels, respectively. The parameters $m^D_{\text{MT200}}$ and $m^A_{\text{MT200}}$ represent the slopes obtained for the donor and acceptor dyes using the MT200, whereas $m^D_{\text{FluoroMax}}$ and $m^A_{\text{FluoroMax}}$ represent the corresponding slopes obtained using the FluoroMax (Horiba) reference measurements.

For determination of the MT200 response, 1 µM solutions of free Cy3 or sCy5 dye in deionized water were measured for 30 s at excitation powers of 1 µW, 2 µW, and 3 µW. Cy3 was excited at 511 nm and sCy5 at 638 nm using pulsed excitation with a repetition rate of 20 MHz. For each excitation power, the mean fluorescence count rate was determined from the corresponding intensity time trace. The signal obtained from deionized water under identical conditions was subtracted as background. The background-corrected mean count rate was plotted against the excitation power

and fitted with a linear function. The resulting slopes were used as $m^D_{\mathrm{MT200}}$ and $m^A_{\mathrm{MT200}}$ for Cy3 and sCy5, respectively.
Reference measurements were performed using a FluoroMax spectrofluorometer (Horiba). Emission spectra were recorded from 550 nm to 800 nm following excitation at 532 nm. The spectrum obtained from deionized water was subtracted as background. The maximum background-corrected fluorescence intensity of each spectrum was plotted against the corresponding dye concentration and fitted with a linear function. The resulting slopes were used as $m^D_{\mathrm{FluoroMax}}$ and $m^A_{\mathrm{FluoroMax}}$, respectively.
The relative detection efficiency was calculated from the four slopes according to Eq. 1, yielding $\eta_{A/D} = 1.13$. Using slopes obtained from multiple measurement points reduced the influence of variations in individual fluorescence intensity measurements.

##### Sample preparation for $QY$ and $R_0$ determination

Single-labeled RNA oligonucleotides carrying either an sCy5 fluorophore at U10 or a Cy3 fluorophore at the 3′-terminal nucleotide G65 were obtained from Ella Biotech:
5'-UGAAGAAAUUCAAAAAAAAAGCUCGGAAUUUGAGCAAAAAAAAAAAACGGUGGUAAAUUCCAUCG-3'
5'-UGAAGAAAUUCAAAAAAAAAGCUCGGAAUUUGAGCAAAAAAAAAAAACGGUGGUAAAUUCCAUCG-3'
The sCy5- and Cy3-labeled nucleotides are highlighted in red and green, respectively. Rhodamine 6G (R6G) and ATTO 647N were used as reference dyes for Cy3 and sCy5, respectively, with literature reference quantum yields of $QY(\mathrm{R6G}) = 0.92$ in ethanol (2) and $QY(\mathrm{ATTO\,647N}) = 0.88$ in ethanol (ATTO-TEC). Refractive indices of $n = 1.36$ for ethanol and $n = 1.33$ for water were used. Four different metal ion conditions were measured:

- $H_2O$
- HEPES with 116 mM $K^+$
- HEPES with 116 mM $K^+$ and 10 mM $Mg^{2+}$
- HEPES with 116 mM $K^+$ and 100 mM $Mg^{2+}$

Samples were prepared in a total volume of 100 µL. RNA concentrations of 0.1 µM and 0.2 µM and reference dye concentrations of 1 µM and 2 µM were used, allowing the quantum yield to be determined independently of concentration from the slope of the corresponding regression line (vide infra).

##### Spectroscopic measurements for $QY$ and $R_0$

Absorption spectra were recorded in 2 nm increments over the following wavelength ranges:

- Cy3/R6G: 450 nm to 600 nm
- sCy5/ATTO 647N: 500 nm to 700 nm

Emission spectra were recorded with an integration time of 100 ms in 2 nm increments over the following wavelength ranges:

- Cy3/R6G: 520 nm to 675 nm, excitation at 500 nm

- sCy5/ATTO 647N: 598 nm to 775 nm, excitation at 578 nm

For each RNA sample and reference dye, two different concentrations were measured. This enabled linear regression of the integrated fluorescence emission intensity against the absorbance at the excitation wavelength, allowing the quantum yield to be determined from the slope of the regression line independently of the absolute sample concentration.

##### Quantum yield calculation

The fluorescence quantum yield ($QY$) was calculated according to

$$QY = QY_{\mathrm{R}} \cdot \frac{I}{I_{\mathrm{R}}} \cdot \frac{OD_{\mathrm{R}}}{OD} \cdot \frac{n^2}{n_{\mathrm{R}}^2} = QY_{\mathrm{R}} \cdot \frac{m}{m_{\mathrm{R}}} \cdot \frac{n^2}{n_{\mathrm{R}}^2}. \tag{2}$$

Here, $m$ and $m_{\mathrm{R}}$ denote the slopes obtained from linear regression of the integrated fluorescence emission intensity against the absorbance at the excitation wavelength for the sample and reference dye, respectively. The refractive indices of the corresponding solvents are denoted by $n$ and $n_{\mathrm{R}}$. The slopes were determined from the two concentrations measured for each sample and reference dye (3, 4).

##### Förster radius $R_0$

The Förster radius $R_0$ was determined experimentally from the spectral overlap integral $J$ according to

$$J = \int j(\lambda)\,\mathrm{d}\lambda, \qquad j(\lambda) = F_{\mathrm{D}}(\lambda) \cdot \varepsilon_{\mathrm{A}}(\lambda) \cdot \lambda^4. \tag{3}$$

The integration range was defined by the spectral overlap between the donor emission and acceptor absorbance spectra. For sCy5, a maximum extinction coefficient of $\varepsilon_{\mathrm{max}} = 250\,000\,\mathrm{M}^{-1}\,\mathrm{cm}^{-1}$ was used. The resulting spectral overlap function $j(\lambda)$ is shown in Figure S10.
Using $J$ in units of $\mathrm{M}^{-1}\,\mathrm{cm}^{-1}\,\mathrm{nm}^4$, the orientation factor $\kappa^2$, the refractive index $n$, and the donor fluorescence quantum yield $QY_{\mathrm{D}}$, the Förster radius was calculated according to (5, 6)

$$R_0\,(\text{Å}) = 0.2108 \left( \frac{\kappa^2 QY_{\mathrm{D}}}{n^4} J \right)^{1/6}. \tag{4}$$

##### smFRET measurements of the IBA-purchased construct

The smFRET data of the IBA-purchased KL–$\mathrm{TL_{GAAA}}$ construct were previously reported by Weber et al. (7) and are included here as a reference for the unbound conformational ensemble. Measurements were performed on a home-built confocal microscope using pulsed-overlaid excitation (POE) under the conditions described previously (7). Briefly, RNA was measured freely diffusing in solution in 50 mM Tris-HCl, pH 7.5, containing 116 mM KCl containing $MgCl_2$ concentrations ranging from 0 mM to 10 mM at room temperature. Cy3 and sCy5 were excited at 532 nm and 638 nm, respectively, and fluorescence was detected in donor and acceptor channels using 582/64 nm and 690/70 nm band-pass filters.

Bursts were identified using a two-color all-photon burst search with a minimum total photon count of 40. Double-labeled molecules were selected using $N_{\mathrm{A,A}} > 20$ and $S > 0.2$, and FRET histograms were corrected according to established procedures (5, 8, 9). The corresponding correction factors are summarized elsewhere (7).

##### smFRET measurements of the Ella Biotech-purchased construct

The smFRET data of the Cy3/sCy5-labeled RNA KL–$TL_{GAAA}$ construct purchased from Ella Biotech are also included here as a reference for the unbound conformational ensemble. Measurements were performed on a commercial confocal fluorescence microscope (MicroTime 200, PicoQuant) using pulsed interleaved excitation (PIE). The PIE sequence was operated at 20 MHz using pulsed 511 nm and 638 nm excitation at 20 $\mu$W and 10 $\mu$W, respectively, measured at the sample position after the objective. Briefly, RNA was measured freely diffusing in standard buffer (20 mM HEPES, pH 7.5, 100 mM KCl) containing $MgCl_2$ concentrations ranging from 0 mM to 100 mM at room temperature. Cy3 and sCy5 were excited at 511 nm and 638 nm, respectively, and fluorescence was detected in donor and acceptor channels using 582/64 nm and 690/70 nm band-pass filters.

Bursts were identified using a two-color all-photon burst search with a minimum total photon count of 50. Bursts were retained for corrected stoichiometries of $-0.10 \leq S \leq 1.10$ and ALEX-2CDE values $\leq 15$ (10), and FRET histograms were corrected according to established procedures (5, 8, 9). The corresponding correction factors are summarized in Supplementary Table S1. The Förster radius of the Cy3/sCy5 pair is shown in Supplementary Table S2.

#### Additional ensemble fluorescence acquisition parameters

For measurements following donor excitation, an excitation wavelength of 530 nm was used, and the donor and acceptor fluorescence intensities were evaluated at 595 nm and 670 nm, respectively. Direct acceptor excitation was recorded using an excitation wavelength of 630 nm. Measurements were performed using an integration time of 100 ms. Ensemble FRET efficiencies were determined from the corrected donor and acceptor fluorescence intensities according to (11).

#### Fluorescence anisotropy and lifetime

##### Sample preparation for lifetime measurements

Three buffer conditions relevant to the analysis of the bound KL–$TL_{GAAA}$ state were investigated:

- HEPES with 116 mM $K^+$
- HEPES with 116 mM $K^+$ and 10 mM $Mg^{2+}$
- HEPES with 116 mM $K^+$ and 100 mM $Mg^{2+}$

Samples were prepared in a total volume of 100 µL at an RNA concentration of 1 µM. All samples contained 20 mM HEPES adjusted to pH 7.5.

##### Anisotropy-based calculation of the cone semiangle ($\theta_0$)

To calculate the cone semiangle $\theta_0$, the Lipari-Szabo wobbling-in-a-cone model was used (12). The generalized orientational order parameter ($S$) was related to $\theta_0$ according to

$$S = \frac{1}{2}\cos(\theta_0)\left[1+\cos(\theta_0)\right]. \tag{5}$$

The order parameter was determined from the fit parameters of the global rotational model applied to the fluorescence anisotropy decays. Specifically, $S$ was calculated from $r_0$ and $r_\infty$ as

$$S = \sqrt{\frac{r_\infty}{r_0}}. \tag{6}$$

The cone semiangle was then obtained by solving the wobbling-in-a-cone relation for $\theta_0$:

$$\theta_0 = \arccos\left[\frac{\sqrt{1+8S}-1}{2}\right]. \tag{7}$$

##### Anisotropy-based calculation of $R_h$

Time-resolved fluorescence anisotropy decays were fitted using the local-global rotational tumbling model (13). The slow rotational correlation time obtained from the fit, $\tau_{\mathrm{global}}$, was assigned to the global tumbling of the dye-labeled KLTL complex. An effective rotational hydrodynamic radius, $R_{\mathrm{h,rot}}$, was calculated from $\tau_{\mathrm{global}}$ using the Stokes-Einstein-Debye relation for rotational diffusion:

$$\tau_{\mathrm{global}} = \frac{4\pi\eta R_{\mathrm{h,rot}}^3}{3k_{\mathrm{B}}T}, \tag{8}$$

which was rearranged to

$$R_{\mathrm{h,rot}} = \left(\frac{3k_{\mathrm{B}}T\tau_{\mathrm{global}}}{4\pi\eta}\right)^{1/3}. \tag{9}$$

Here, $k_{\mathrm{B}}$ is the Boltzmann constant, $T$ is the absolute temperature, and $\eta$ is the dynamic viscosity of the solvent under the corresponding experimental or simulation conditions. The conversion was applied independently to the $\tau_{\mathrm{global}}$ values obtained for Cy3 and sCy5 from the experimental anisotropy decays and from the restrained and unrestrained simulations. The resulting radius represents an effective rotational hydrodynamic radius under the assumption of approximately isotropic rotational diffusion.

##### Structure-based calculation of $R_h$ using HullRad

A structure-based rotational hydrodynamic radius was calculated from the bound KLTL structure using HullRad version 10.1 (14). The atomic coordinates were supplied to HullRad in PDB format. HullRad generated a reduced representation of the molecular structure and calculated its convex-hull surface area, volume, maximum dimension, and shape-dependent frictional properties. An empirical hydration layer and a shape correction were then applied to determine the effective rotational radius reported by HullRad as `R(Rotation)`.
The rotational diffusion coefficient ($D_{\mathrm{r}}$) was calculated according to

$$D_{\mathrm{r}} = \frac{k_{\mathrm{B}}T}{8\pi\eta R_{\mathrm{h,rot}}^3}, \tag{10}$$

and the corresponding second-rank rotational correlation time was obtained from

$$\tau_{\mathrm{global}} = \frac{1}{6D_{\mathrm{r}}} = \frac{4\pi\eta R_{\mathrm{h,rot}}^3}{3k_{\mathrm{B}}T}. \tag{11}$$

The HullRad calculation used the solvent parameters implemented in the program, corresponding to water at 293.15 K with a dynamic viscosity of 1.001 67 mPa s. The value reported as `R(Rotation)` was used for comparison with the effective rotational hydrodynamic radii derived from the anisotropy measurements and simulations. The translational radius reported by HullRad as `R(Translation)` was not used for this comparison.

#### MD Simulations

All molecular dynamics simulations were performed with GROMACS release 2025.3 and were set up and executed through the `MDSimulation` class of the FAMP pipeline (15). FAMP creates the working directory structure, copies the force field directory, the `.mdp` parameter files and the input structure into it, and executes the individual GROMACS commands sequentially via the Python `subprocess` module. The user-defined input for MD simulations with FAMP is reduced to simulation time, temperature, $Mg^{2+}$ concentration, solute–box-edge distance, water model and the distance-restraint flag.

##### Forcefield

The RNA was described by the AMBER force field (16) with the bsc0 (17) and $\chi_{\mathrm{OL3}}$ (18, 19) corrections. More recent RNA force field parameterizations are available (20); $\chi_{\mathrm{OL3}}$ was retained here because it is broadly transferable across RNA structural motifs, whereas newer developments tend to be optimized towards, and potentially over-parameterized for, individual motifs. For a construct combining a kissing loop and a GAAA tetraloop–receptor contact, a generally applicable parameter set was considered preferable.
Ion parameters for $K^+$ and $Cl^-$ were taken from Joung and Cheatham (21), and $Mg^{2+}$ was described with the parameters of Allnér, Nilsson and Villa (22). Water was modelled with TIP4P-Ew (23). Force field, ion and water parameters were taken from the distribution provided with FRETlabel (13, 24).
The acceptor sCy5, including its linker, was described by AMBER-DYES (25, 26) as distributed with FRETlabel (13). For the donor Cy3, no parameter set matching the linker chemistry of the labelled construct was available. A fragment was therefore built from the C3W structure by removing the sulfo groups and the linker, rebuilding the linker to match the attachment chemistry specified by the manufacturer (Ella Biotech), and capping the termini to give a net charge of $-1$. The fragment geometry was optimized at the B3LYP/6-31G level and the electrostatic potential computed at the HF/6-31G level with Gaussian. Partial charges were obtained from a two-stage RESP fit with AmberTools23 (27); in contrast to the fragments distributed with FRETlabel (24), no group constraints were applied to the capping groups. Only partial charges were derived here; bonded and Lennard-Jones parameters were taken from AMBER-DYES, with terms for the rebuilt linker atoms assigned by analogy to existing AMBER parameters using ACPYPE (28). The resulting fragment is included in the force field directory provided with FAMP (15).

##### Solvation

The *in silico* labelled starting structure was converted to GROMACS format and topology with `pdb2gmx`, using the force field provided by the pipeline. A dodecahedral box was defined with `editconf` at a minimum distance of 1.25 nm between the solute and the box edge, and filled with explicit TIP4P-Ew water (23) using `solvate`.
Ions were added by randomly replacing water molecules. $Mg^{2+}$ was set to 20 mM or 100 mM; at the experimental concentration of 10 mM (29), a box of this size would contain only two to four ions, which is insufficient to describe the ionic atmosphere (30). Each system was solvated and ionised independently; the resulting compositions and $Mg^{2+}$ concentrations of 19.8 and 101.8 mM are listed in Table S5.

#### MD run

Energy minimization was performed with `mdrun` under positional restraints on the RNA heavy atoms, allowing the water molecules to relax around the solute. Equilibration was carried out in two steps of 300 ps each. In the canonical ensemble (NVT), the temperature was set to 298 K using the velocity-rescale thermostat (31). In the isothermal–isobaric ensemble (NPT), the pressure was set to 1 bar using the Parrinello–Rahman barostat (32) at the same temperature.
Production runs of 1000 ns were performed with the leap-frog integrator (33) at an integration time step of 2 fs. Bonds were constrained with the LINCS algorithm (34). Non-bonded interactions were treated with the Verlet cut-off scheme at a cut-off length of 1.0 nm, and long-range electrostatics with the particle mesh Ewald method (35).
Four production runs were performed, combining the two $Mg^{2+}$ concentrations with the presence or absence of distance restraints (Table S5). All four systems were prepared from the same labelled starting structure and are identical in every parameter other than those listed.
In the restrained simulations, the docking of the GAAA tetraloop onto its receptor was maintained by six pairwise distance restraints (Table S4). The atom pairs and the corresponding distances were taken from the reference structure PDB 3JCT; polar contacts were identified in PyMOL using the `distance` command in mode 2. Restraints were implemented as GROMACS distance restraints (`[ distance_restraints ]`, type 1) with a linear/harmonic potential defined by the three distances $r_0$, $r_1$ and $r_2$
The lower bound $r_0$ was set to zero for all pairs, $r_1$ was set to the atom-atom distance measured in 3JCT and $r_2$ to $r_1$+0.20 nm. The restraining force is capped at $k_{dr}(r_2-r_1)$. With a force constant of 1000 kJ mol$^{-1}$ nm$^{-2}$ and a weighting factor of 0.25 applied to every pair, the effective force constant is $k_{dr}$ = 250 kJ mol$^{-1}$ nm$^{-2}$. Restraints were applied as simple (non-ensemble) restraints without time averaging (`disre = simple`, `disre_tau = 0`) and were applied during the MD long run.

#### *in silico* FRET predictions

The following parameters were used in FRETlabel to calculate the ACVs for each structure and trajectory frame, with condition-specific `cv_fraction` values derived from the corresponding dynamic fluorescence anisotropy measurements at the respective $Mg^{2+}$ concentrations.

```
{
    "Position": {
        "Cy3-65-O3'": {
            "attach_id": 2103,
            "mol_selection": "all",
            "linker_length": 6,
            "linker_width": 3.5,
            "dye_radius1": 8,
            "dye_radius2": 3,
            "dye_radius3": 1.5,
            "cv_fraction": "<cv_fraction>",
            "cv_thickness": 3,
            "use_LabelLib": false,
            "grid_spacing": 1.0,
            "simulation_type": "AV3",
            "state": 1,
            "frame_mdtraj": 0,
            "contour_level_AV": 0,
            "contour_level_CV": 0.7,
            "b_factor": 100,
            "gaussian_resolution": 2,
```

```
            "grid_buffer": 2.0,
            "transparent_AV": true
        },
        "Cy5-10-C5": {
            "attach_id": 307,
            "mol_selection": "all",
            "linker_length": 20,
            "linker_width": 3.5,
            "dye_radius1": 9.5,
            "dye_radius2": 3,
            "dye_radius3": 1.5,
            "cv_fraction": "<cv_fraction>",
            "cv_thickness": 3,
            "use_LabelLib": false,
            "grid_spacing": 1.0,
            "simulation_type": "AV3",
            "state": 1,
            "frame_mdtraj": 0,
            "contour_level_AV": 0,
            "contour_level_CV": 0.7,
            "b_factor": 100,
            "gaussian_resolution": 2,
            "grid_buffer": 2.0,
            "transparent_AV": false
        }
    },
    "Distance": {
        "Cy3-Cy5": {
            "R0": "<R0_value>",
            "n_dist": 1000000
        }
    }
}
```

The following parameters were used for photon sampling with FRETraj across all structure collections to generate both unweighted and weighted FRET distributions, using the corresponding Rkappa files.

```
{
    "dyes": {
        "tauD": 1.4,
        "tauA": 1.12,
        "QD": 0.46,
        "QA": 0.35,
        "etaA": 1,
        "etaD": 0.37,
        "dipole_angle_abs_em": 0
    },
    "sampling": {
        "nbursts": 20000,
        "skipframesatstart": 0,
        "skipframesatend": 0,
        "multiprocessing": true
    },
    "fret": {
        "R0": 61.7,
        "kappasquare": 0.6666,
        "gamma": true,
        "quenching_radius": 1
```

```
    },
    "species": {
        "name": ["all"],
        "unix_pattern_rkappa": ["<unweighted> or <weighted> r_kappa file"],
        "unix_pattern_don_coords": [],
        "unix_pattern_acc_coords": [],
        "probability": [1],
        "n_trajectory_splits": null
    },
    "bursts": {
        "lower_limit": null,
        "upper_limit": null,
        "lambda": null,
        "QY_correction": false,
        "averaging": "trajectory",
        "burst_size_file": "<experimental burst sizes>"
    }
}
```

#### Trajectory processing and structural analysis

All structural analyses of the MD trajectories were performed using the Python package MDAnalysis (36, 37) within the `DataAnalysis` class of FAMP (38). Root mean square deviations (RMSD) and root mean square fluctuations (RMSF) were calculated after aligning the KL domain (nucleotides 24–33 and 53–62) of each trajectory frame to the initial input structure. This alignment isolates fluctuations of the TL and poly(A) linker regions from the overall rotational and translational motion of the construct. Atomic coordinates were extracted frame-wise to calculate the time-dependent inter-dye distances and orientation factors used for the FRET analysis.

#### Inter-dye distance and orientation factor from explicit dyes

Atom identifiers of the dye centers and the atoms defining the transition dipole vectors were determined automatically from the residue names and residue numbers in the `.gro` and `.pdb` files using the `Dye` class of FAMP. The inter-dye distance $R_{\mathrm{DA}}(t)$ was calculated as the Euclidean distance between the central carbon atoms of the polymethine chains of Cy3 (C11) and sCy5 (C32),

$$R_{\mathrm{DA}}(t) = \|\mathbf{r}_{\mathrm{C11}}(t) - \mathbf{r}_{\mathrm{C32}}(t)\| = \sqrt{\sum_{\alpha=x,y,z} [r_{\mathrm{C11},\alpha}(t) - r_{\mathrm{C32},\alpha}(t)]^2}. \tag{S12}$$

The orientation factor $\kappa^2(t)$ was calculated from the normalized donor and acceptor transition dipole vectors according to

$$\kappa^2(t) = [\cos\theta_{\mathrm{T}}(t) - 3\cos\theta_{\mathrm{D}}(t)\cos\theta_{\mathrm{A}}(t)]^2 . \tag{S13}$$

The transition dipole vectors were defined by the vector connecting atoms C2 and C14 of the respective fluorophore. Here, $\cos\theta_{\mathrm{T}}$ denotes the scalar product of the normalized donor and acceptor transition dipole vectors, whereas $\theta_{\mathrm{D}}$ and $\theta_{\mathrm{A}}$ describe the angles between the respective transition dipole vectors and the normalized donor–acceptor connecting vector (39, 40).

#### mACV calculation along the MD trajectories

mACVs were calculated from the MD trajectories without explicit dyes. The reduced `.xtc` trajectories and corresponding PDB structures were loaded with MDTraj (41), and mACVs were calculated every 100 ps using `fretraj.cloud.pipeline_frames()`. Dye attachment atoms were identified using the `Dye` class of FAMP, while the remaining FRETraj parameters were taken from the dye parametrization listed in SI Table 6. The `cv_fraction` was derived from the corresponding experimental fluorescence anisotropy measurements (SI Table 7) to account for the measured dye–RNA stacking contribution (13, 24).
The resulting mACVs were analyzed using `fretraj.cloud.Trajectory()` assuming an isotropic orientation factor of $\kappa^2 = 2/3$. For each frame, the donor–acceptor distance $R_{\mathrm{DA}}(t)$, transfer efficiency $E_{\mathrm{DA}}(t)$, distance between the mean dye positions $R_{\mathrm{MP}}(t)$, and attachment-point distance $R_{\mathrm{AP}}(t)$ were calculated. The resulting data were written to `r_kappa.dat` files and stored as `.pkl` objects for further analysis.

#### Per-frame transfer efficiency

For both dye representations, the transfer efficiency of each trajectory frame was calculated according to

$$E_{\mathrm{DA}}(t) = \frac{1}{1 + \dfrac{R_{\mathrm{DA}}(t)^6 (2/3)}{R_0^6 \kappa^2(t)}}. \tag{S14}$$

Here, $R_0$ is the experimentally determined Förster radius of the Cy3/sCy5 pair (SI Table 6). Since $R_0$ was determined assuming isotropic orientational averaging with $\kappa^2 = 2/3$, the factor $(2/3)/\kappa^2(t)$ accounts for the instantaneous dye orientation in the explicit-dye trajectories. For the mACV representation, $\kappa^2(t) = 2/3$ was assumed for all frames, reducing Eq. (S14) to

$$E_{\mathrm{DA}}(t) = \left[1 + \left(\frac{R_{\mathrm{DA}}(t)}{R_0}\right)^6\right]^{-1}. \tag{S15}$$

#### Supplementary Figures

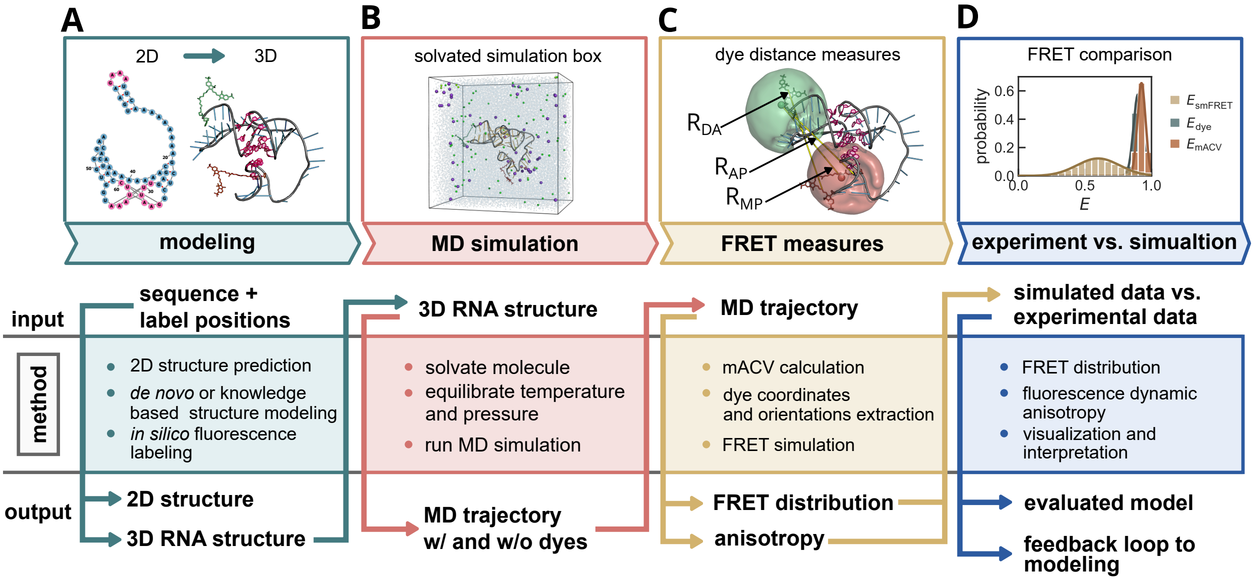


Fig. S1: Extended methods of the FAMP pipeline: **A)** *de novo* modelling is performed using Rosetta FARFAR2. A knowledge-based approach. The generated tertiary structure model is in silico labelled with fluorescent dyes. **B)** dynamics of the RNA and dyes are simulated using MD simulations. **C)** dye coordinates are extracted from MD trajectories, and FRET distributions are simulated using either MACV or explicit dyes. This generates FRET distributions, anisotropy decays, and other intra-molecular distances. **D)** simulated FRET data is compared with fully corrected experimental FRET data.

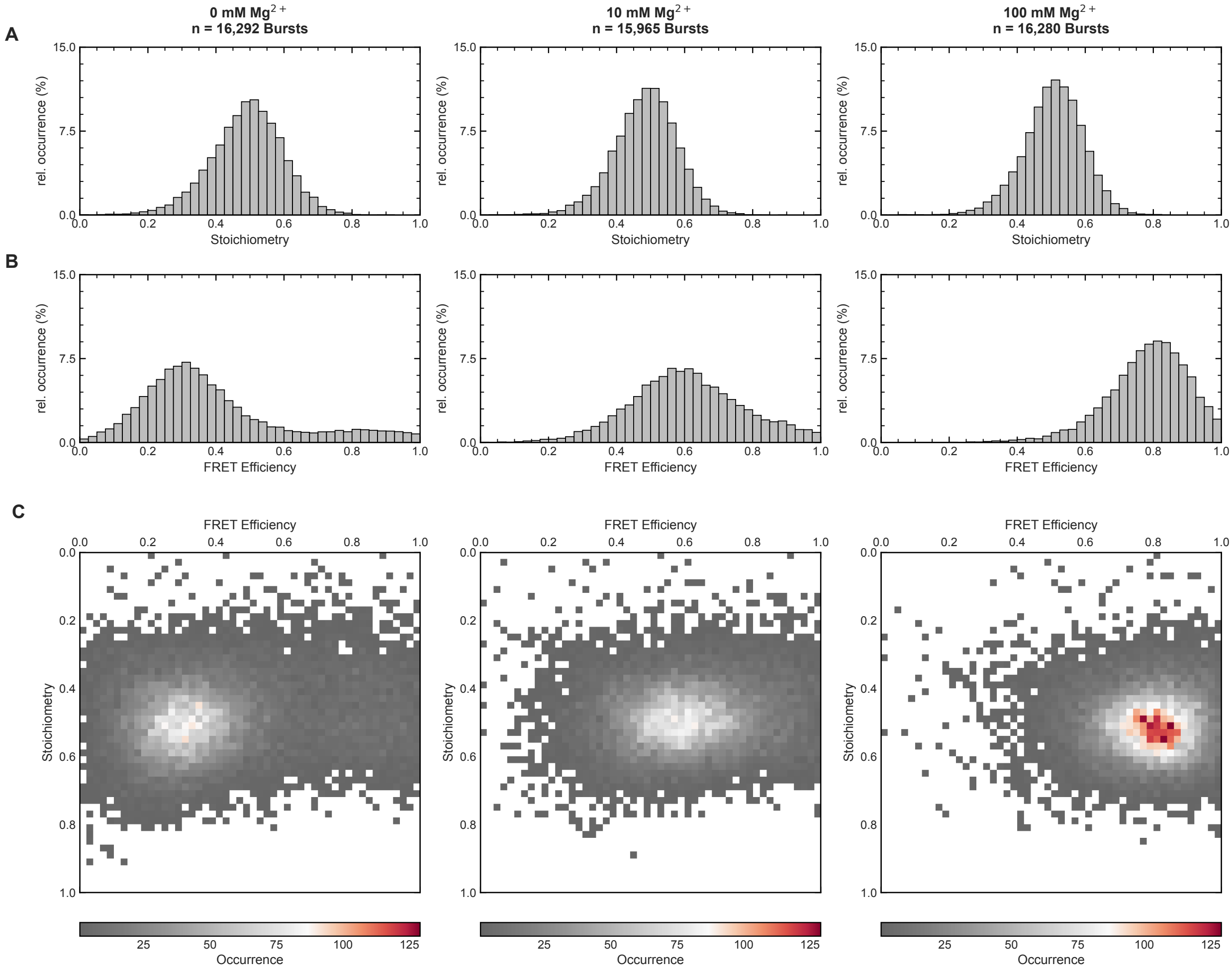


Fig. S2: Freely diffusing, doubly labeled (Cy3/sCy5) RNA was measured in 20 mM HEPES, pH 7.5, 116 mM KCl at RT in the presence of 0 mM (left), 10 mM (middle) and 100 mM (right) $MgCl_2$ using a MicroTime 200 confocal fluorescence microscope (PicoQuant) operated in pulsed interleaved excitation (PIE) mode in HybriWell microfluidic chambers (Grace Biolabs). Data were fully corrected for background, spectral bleed-through, direct excitation, detection efficiency and quantum yield, and donor-only species were excluded by molecular sorting. **(A)** Stoichiometry histograms, **(B)** FRET efficiency histograms and **(C)** two-dimensional stoichiometry–FRET histograms.

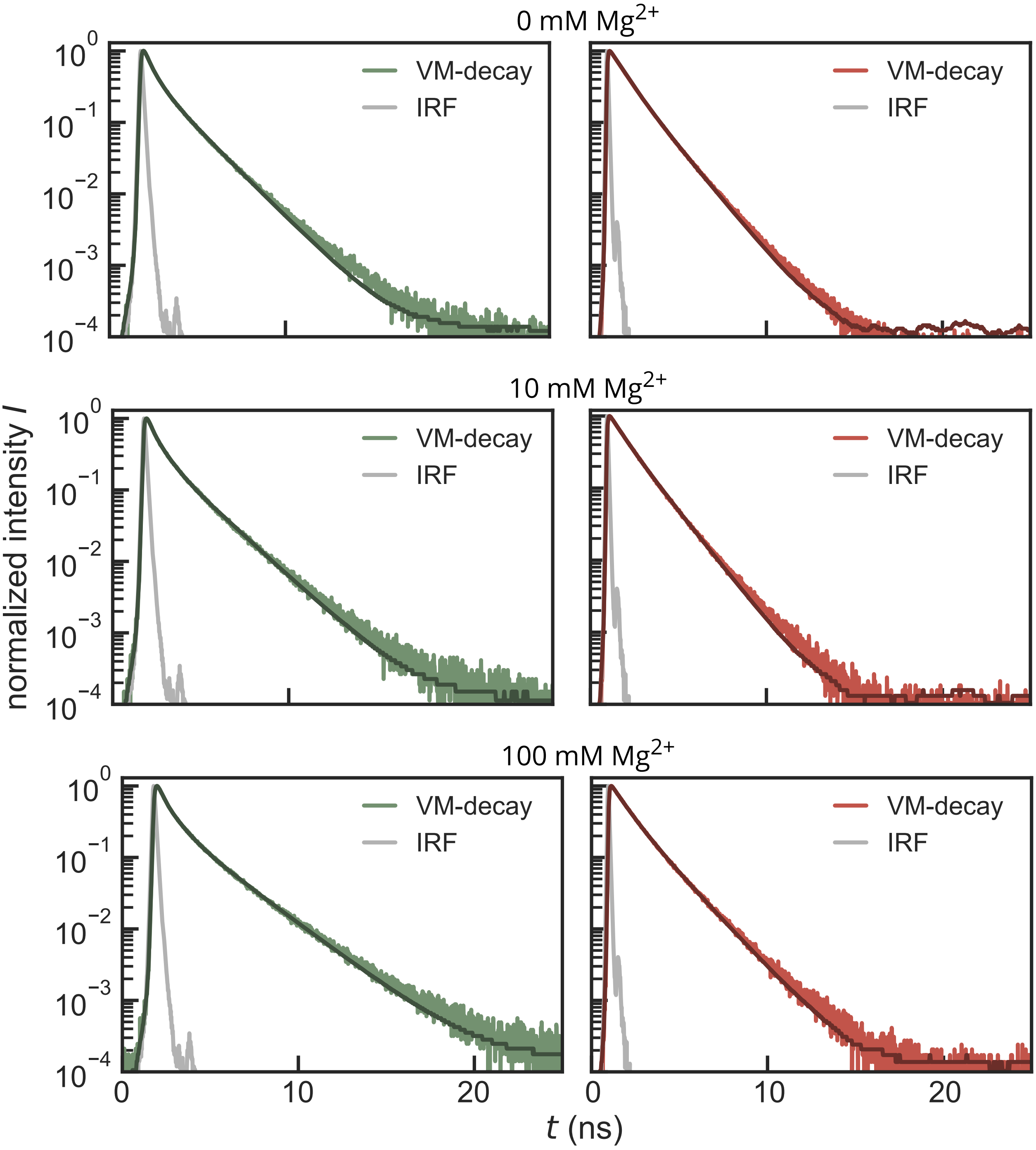


Fig. S3: Fluorescence lifetimes decays of Cy3 (green) and Cy5 (red) bound to the KL-TLGAAA construct, including the instrument response function (IRF).

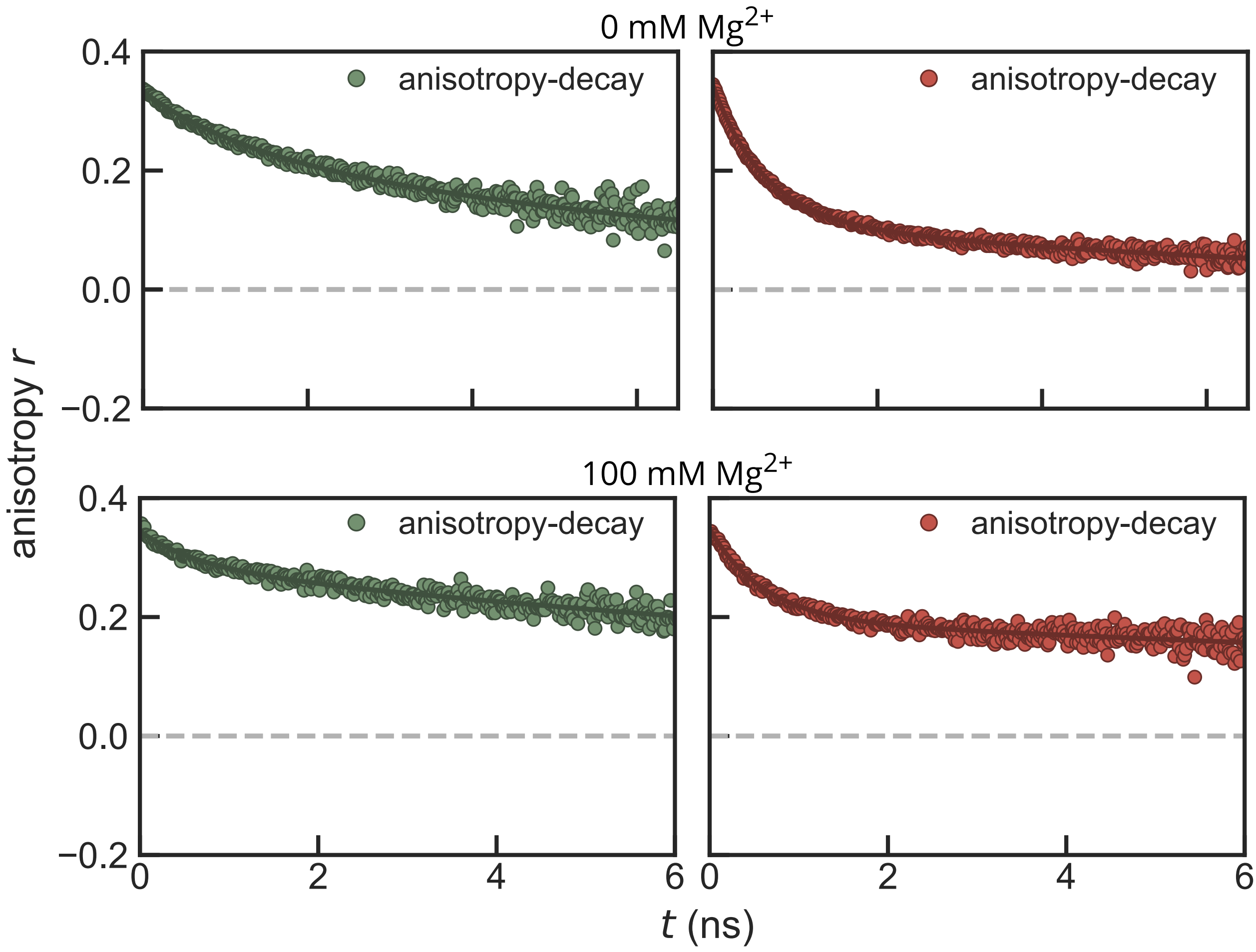


Fig. S4: Anisotropy decays of Cy3 (green) and Cy5 (red) bound to the KL-TLGAAA construct, fitted with the local–global rotation-in-acone model.

**A**

$$V_{dr}(r_{ij}) = \begin{cases} \frac{1}{2}k_{dr}(r_{ij} - r_{low})^2 & \text{for} \quad r_{ij} < r_{low} \\ 0 & \text{for} \quad r_{low} \leq r_{ij} < r_{up\,1} \\ \frac{1}{2}k_{dr}(r_{ij} - r_{up\,1})^2 & \text{for} \quad r_{up\,1} \leq r_{ij} < r_{up\,2} \\ \frac{1}{2}k_{dr}(r_{up\,2} - r_{up\,1})(2r_{ij} - r_{up\,2} - r_{up\,1}) & \text{for} \quad r_{up\,2} \leq r_{ij} \end{cases}$$

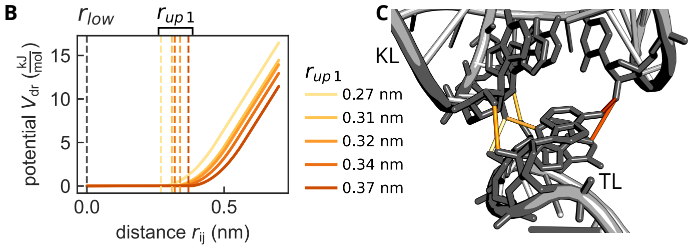


Fig. S5: Distance restraints applied in the restrained simulations. (**A**) GROMACS type-1 distance restraint potential $V_{dr}(r_{ij})$: zero between $r_{\text{low}}$ and $r_{\text{up1}}$, harmonic between $r_{\text{up1}}$ and $r_{\text{up2}}$, and linear beyond, capping the restraining force at $k_{dr}(r_{\text{up2}} - r_{\text{up1}})$. (**B**) Potentials for the six restrained atom pairs. $r_{\text{low}}$ was set to zero, $r_{\text{up1}}$ to the distance measured in the reference structure (PDB 3JCT) and $r_{\text{up2}}$ to $r_{\text{up1}} + 0.20\,\text{nm}$, with $k_{dr} = 250\,\text{kJ}\,\text{mol}^{-1}\,\text{nm}^{-2}$. (**C**) The restraints connect residues A6–A8 of the GAAA tetraloop (TL) to U31 and A55–A57 of the kissing loop (KL) and correspond to the A-minor interactions of the tertiary contact.

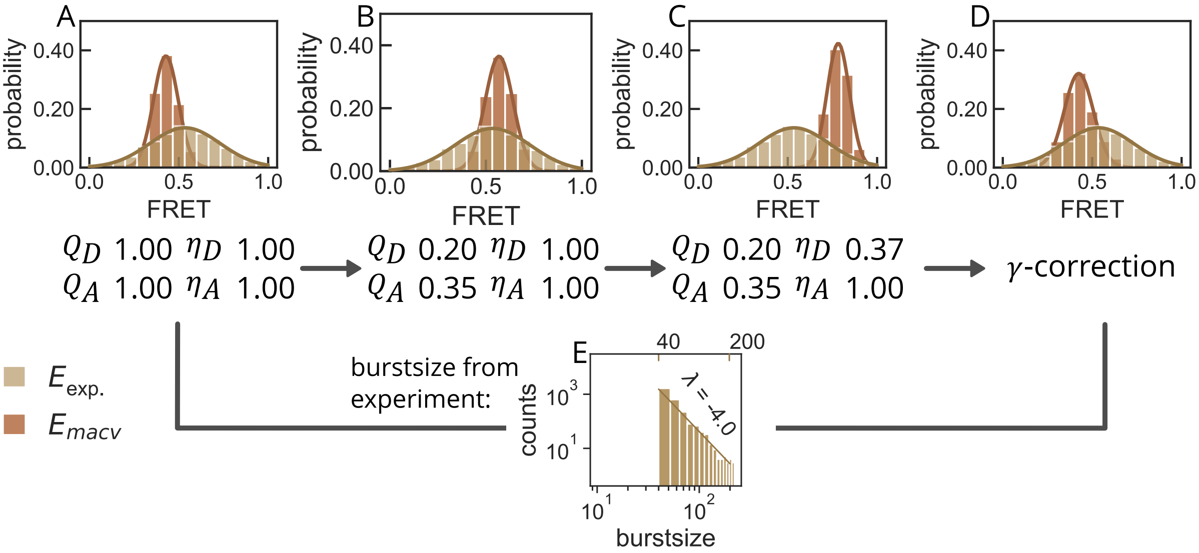


Fig. S6: Influence of experimental parameters on the FRET distribution derived from the unrestrained mACV simulation trajectory. Experimental FRET distributions ($E_{\text{exp.}}$, beige) are compared with distributions calculated from the mACV trajectory ($E_{\text{mACV}}$, orange). **(A)** Simulated FRET distribution assuming ideal experimental conditions, with donor and acceptor quantum yields ($Q_D$, $Q_A$) and detection efficiencies ($\eta_D$, $\eta_A$) set to 1. **(B)** Incorporation of the experimentally determined dye quantum yields ($Q_D = 0.20$, $Q_A = 0.35$) shifts the simulated FRET distribution. **(C)** Additional consideration of the donor detection efficiency ($\eta_D = 0.37$, $\eta_A = 1.00$) further alters the apparent FRET efficiency. **(D)** Subsequent $\gamma$-correction compensates for these experimental effects and restores the mean FRET value obtained under ideal conditions. **(E)** Burst sizes for the simulated data were sampled from the experimental burst-size distribution in the range of 40–200 photons per burst, which follows an approximately power-law behavior with an exponent of 4.0. Incorporating the experimental burst-size distribution reproduces the photon-counting noise and broadening of the simulated FRET distribution, enabling a more direct comparison with the experiment.

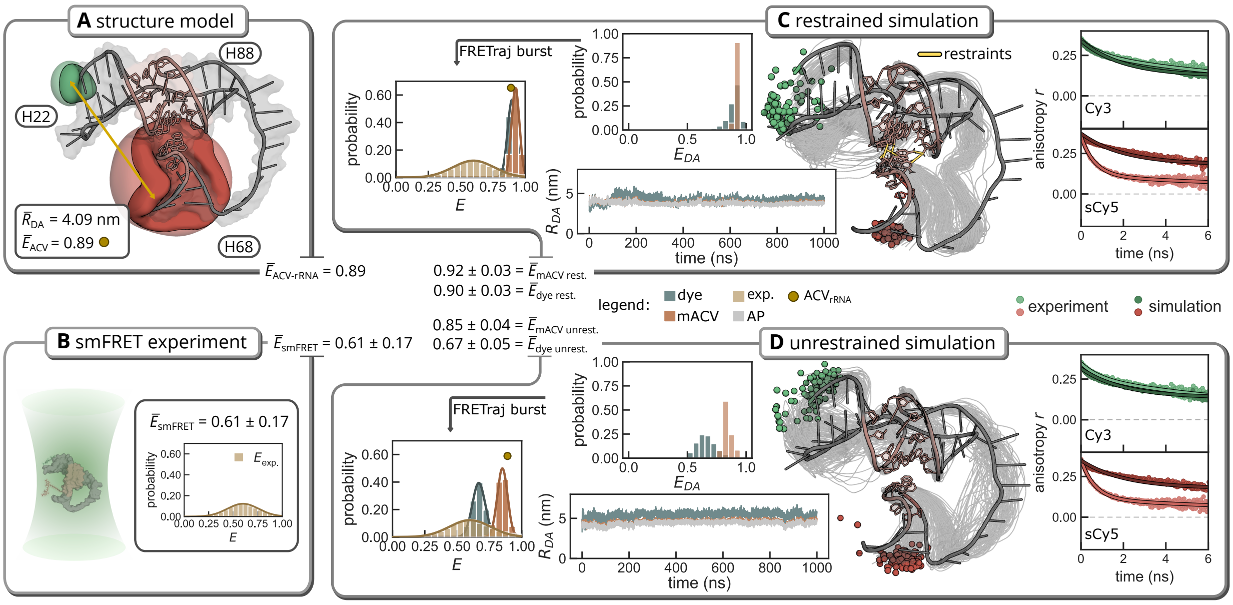


Fig. S7: Comparison of experimental smFRET data with a single structure estimate and restrained and unrestrained MD structure collections. **(A)** Modeled bound state of KL–$TL_{GAAA}$ showing the predicted ACVs, the mean donor–acceptor distance $R_{DA}$, and the corresponding $E_{ACV}$ value. **(B)** Experimental smFRET distribution at an $Mg^{2+}$ concentration of 10 mmol $L^{-1}$, including the corresponding mean and standard deviation. Both **(C)** restrained MD simulation and **(D)** unrestrained MD simulation show the mean C-atoms of the respective dyes Cy3 and sCy5, together with the dye-, attachment-point-, and mACV-derived $R_{DA}$ distances shown in the lower left. The distance-derived $E_{DA}$ distribution is shown in the upper left, whereas the photon-burst-derived FRET histogram, which enables direct comparison with the experiment, is shown on the left for each simulation. Experimental anisotropy measurements and comparison to the all-atom dye simulations are shown on the right for both simulations.

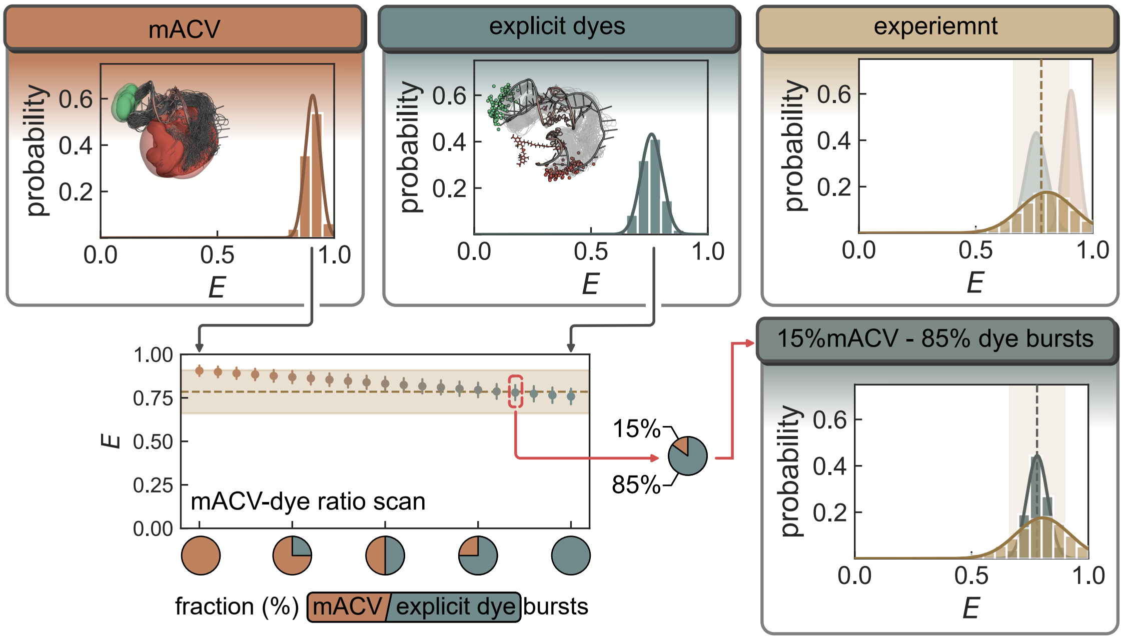


Fig. S8: Merging of burst distributions of the bound state. In the bound state, burst simulations using the mACV model (left) and explicit dyes (middle) yield two narrow distributions, each corresponding to one of the two states contained in the experimental distribution (right). To identify the mixing ratio that reproduces the experimental mean, bursts from both simulations were merged into a joint ensemble while the mACV fraction was varied stepwise (ratio scan, bottom left). Best agreement is obtained for 15% mACV and 85% explicit-dye bursts (bottom right), where the mean of the merged distribution matches the measured one within experimental error.

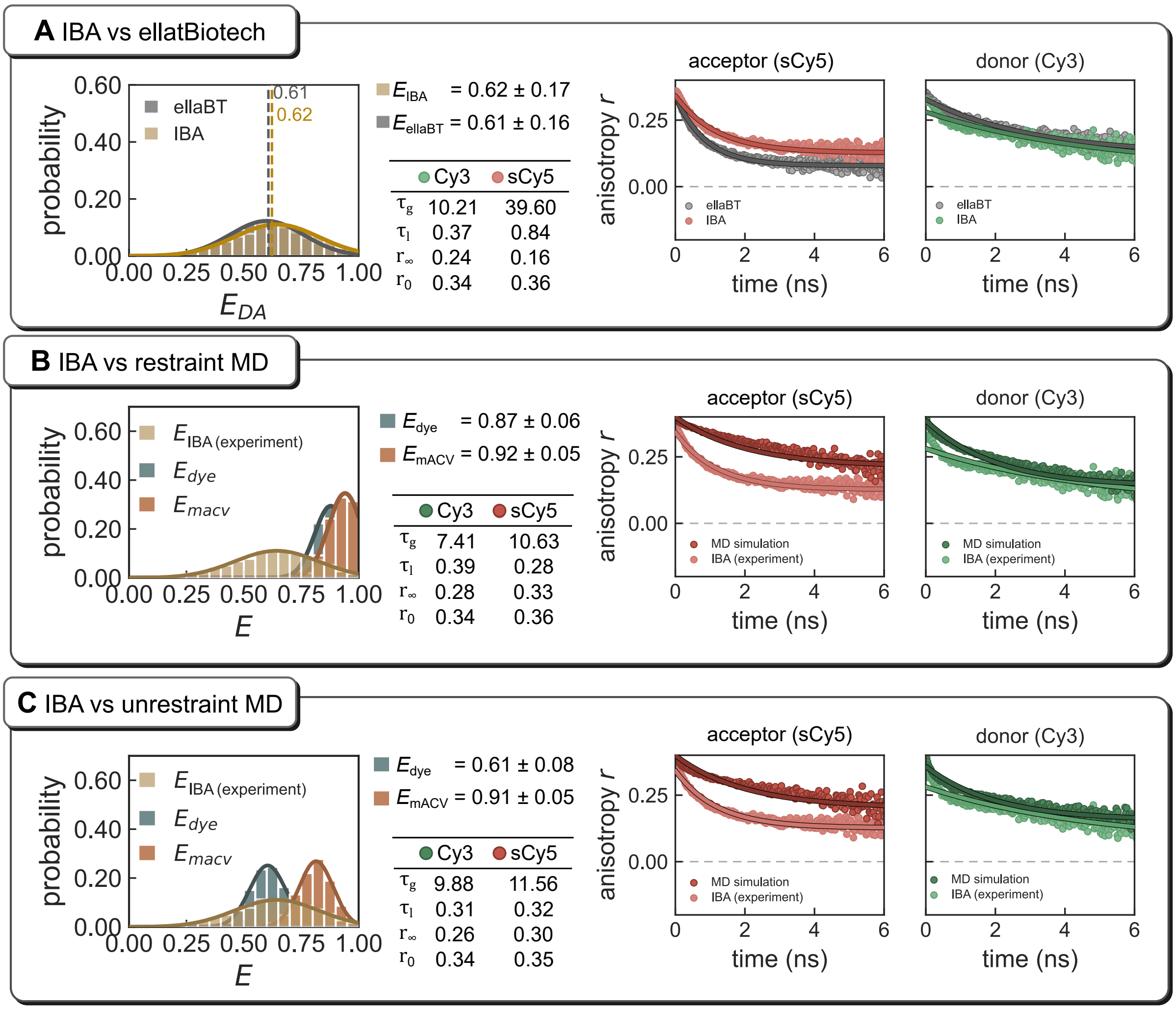


Fig. S9: Comparison of the IBA Lifesciences construct with the Ella Biotech construct and MD simulation results. The same RNA construct obtained from a second supplier (IBA Lifesciences) was measured and analysed independently; all correction factors (quantum yields, detection efficiencies and fluorescence lifetimes) were re-determined for this sample at $10\,mol\,L^{-1}$ $Mg^{2+}$ with $K^{+}$ background. **(A)** smFRET efficiency histograms (left) and time-resolved anisotropy decays of the acceptor sCy5 and the donor Cy3 (right) for the IBA and the Ella Biotech sample. Both constructs yield essentially identical mean transfer efficiencies ($E_{\mathrm{IBA}} = 0.62 \pm 0.17$ vs. $E_{\mathrm{EllaBT}} = 0.61 \pm 0.16$) and comparable rotational dynamics, confirming that the reference data used in the main text are independent of the supplier. **(B, C)** The IBA data set compared with burst simulations from the restrained (B) and unrestrained (C) MD ensembles, using both the explicit-dye and the mACV model, together with the corresponding simulated and experimental anisotropy decays. Tables list the global rotational correlation time $\tau_{\mathrm{g}}$, the fast local correlation time $\tau_{\mathrm{l}}$ (both in ns), the residual anisotropy $r_{\infty}$ and the fundamental anisotropy $r_0$. As for the Ella Biotech sample, the mACV model overestimates the transfer efficiency in both ensembles, whereas the explicit-dye simulation of the unrestrained ensemble reproduces the experimental mean; the agreement between simulation and experiment is therefore of the same quality for both constructs.

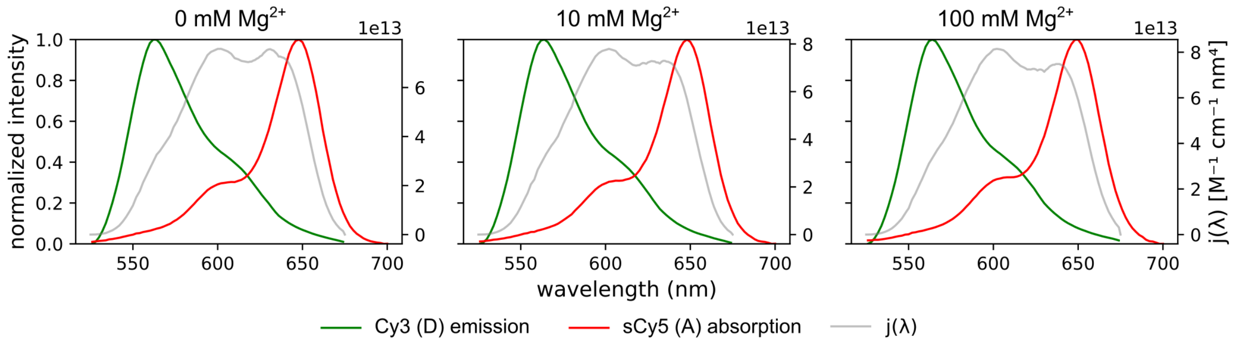


Fig. S10: Spectral overlap integral J for different [$Mg^{2+}$] (from left to right): 0 mM, 10 mM and 100 mM. Each plot shows the normalized donor fluorescence (green) and acceptor absorbance (red) spectra (left axis) and the spectral overlap density j($\lambda$) (gray, right axis)

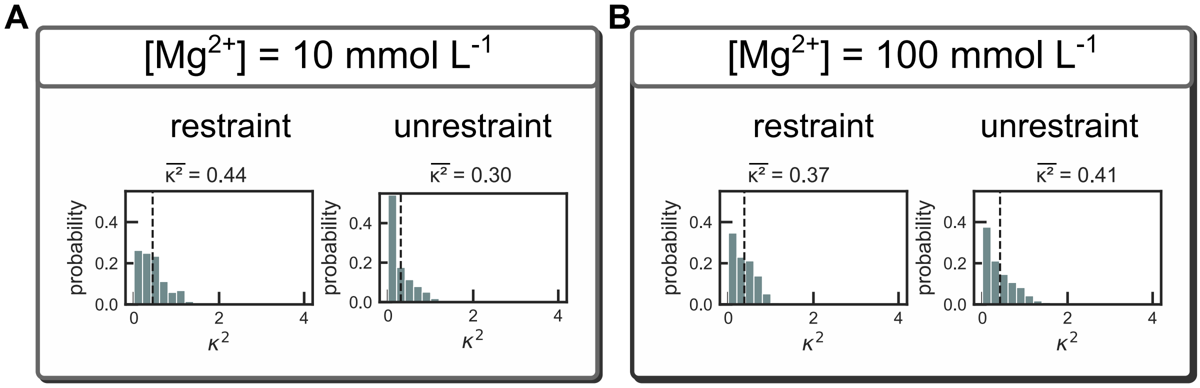


Fig. S11: $\kappa^2$ distributions derived from explicit dye dipole atom coordinates of the restraint and unrestraint simulation of both 10 **(A)** and 100 **(B)** $mmol\,L^{-1}$ $Mg^{+2}$ conditions. All mean $\kappa^2$ showing smaller values than 2/3

#### Supplementary Tables

Table S1: FRET correction factors for spectral crosstalk including bleed-through ($bt$) and direct excitation ($dE$), as well as the detection efficiency ($\eta$) correction factor for detection efficiency.

| **Category** | **Correction factor** | **ellaBiotech** | | **IBA** | |
|---|---|---|---|---|---|
| | | **Donor** | **Acceptor** | **Donor** | **Acceptor** |
| Bleed-through | $bt$ | 8.6% | 0.2% | 9.1% | 0.3% |
| Direct excitation | $dE$ | 0.0% | 3.8% | 0.5% | 4.2% |
| Detection efficiency | $\eta$ | $\eta_A/\eta_D = 1.13$ | | $\eta_A/\eta_D = 2.72$ | |

Table S2: Quantum yield ($Q_{D/A}$), fluorescence lifetimes ($\tau_{D/A}$) and Förster-radius ($R_0$) for all shown $Mg^{2+}$-concentration.

| $c_{Mg^{2+}}$ (mM) | $Q_D$ | $Q_A$ | $\tau_D$ (ns) | $\tau_A$ (ns) | $R_0$ (nm) |
|---|---|---|---|---|---|
| 0 | 0.65 | 0.51 | 1.3 | 1.3 | 6.53 |
| 10 | 0.43 | 0.35 | 1.4 | 1.3 | 6.10 |
| 100 | 0.46 | 0.37 | 1.67 | 1.42 | 6.23 |

Table S3: Fit parameters of the fluorescence lifetime for donor-only (Cy3) and acceptor-only (Cy5) labeled samples at different $Mg^{2+}$ concentrations (13).

| Dye | $[Mg^{2+}]$ (mM) | $\tau_1$ (ns) | $\tau_2$ (ns) | $\tau_3$ (ns) | $\tau_{av}$ (ns) | $a_1$ | $a_2$ | $a_3$ |
|---|---|---|---|---|---|---|---|---|
| Cy3 | 0 | $0.139 \pm 0.002$ | $0.553 \pm 0.004$ | $1.762 \pm 0.004$ | $1.331 \pm 0.004$ | 0.32 | 0.39 | 0.29 |
| sCy5 | 0 | $0.790 \pm 0.007$ | $1.490 \pm 0.007$ | – | $1.280 \pm 0.006$ | 0.45 | 0.55 | – |
| Cy3 | 10 | $0.178 \pm 0.002$ | $0.677 \pm 0.008$ | $1.951 \pm 0.009$ | $1.401 \pm 0.007$ | 0.37 | 0.38 | 0.25 |
| sCy5 | 10 | $0.727 \pm 0.009$ | $1.431 \pm 0.006$ | – | $1.291 \pm 0.005$ | 0.33 | 0.67 | – |
| Cy3 | 100 | $0.222 \pm 0.002$ | $0.803 \pm 0.008$ | $2.386 \pm 0.011$ | $1.672 \pm 0.008$ | 0.37 | 0.40 | 0.23 |
| sCy5 | 100 | $0.810 \pm 0.007$ | $1.671 \pm 0.008$ | – | $1.416 \pm 0.007$ | 0.47 | 0.53 | – |

Table S4: Distance restraints applied in the restrained simulations. Atoms $i$ and $j$ are given as residue type, number and PDB atom name; residue numbering follows the model construct. $r_0$, $r_1$ and $r_2$ define the flat-bottomed potential: no force for $r_0 \leq r \leq r_1$, harmonic below $r_0$ and between $r_1$ and $r_2$, linear above $r_2$. All restraints are of GROMACS type 1 with a weighting factor of 0.25 scaling the global force constant `disre_fc`.

| Index | Atom $i$ | Atom $j$ | $r_0$ (nm) | $r_1$ (nm) | $r_2$ (nm) | Interaction |
|---|---|---|---|---|---|---|
| 0 | A6 O2′ | A57 O2′ | 0.00 | 0.32 | 0.52 | A-minor |
| 1 | A6 N6 | U31 O2′ | 0.00 | 0.37 | 0.57 | A-minor |
| 2 | A7 N6 | U31 O2′ | 0.00 | 0.34 | 0.54 | A-minor |
| 3 | A7 N3 | A56 O2′ | 0.00 | 0.31 | 0.51 | A-minor |
| 4 | A7 O2′ | A56 O2′ | 0.00 | 0.27 | 0.47 | A-minor |
| 5 | A8 O2′ | A55 O2′ | 0.00 | 0.31 | 0.51 | A-minor |

Table S5: Overview of the four MD runs. All systems were prepared from the same *in silico* labelled starting structure and differ only in the $Mg^{2+}$ concentration and the presence of distance restraints on the tertiary contacts. Each system was solvated and ionised independently; the restrained and unrestrained runs of a given $Mg^{2+}$ concentration yielded identical compositions.

| Run | $[Mg^{2+}]$ (mM) | Restraints | Length (ns) | Atoms | Box volume ($nm^3$) | $n(K^+)$ | $n(Mg^{2+})$ |
|---|---|---|---|---|---|---|---|
| 1 | 20 | yes | 1000 | 88 932 | 672.3 | 66 | 8 |
| 2 | 100 | yes | 1000 | 88 635 | 668.7 | 66 | 41 |
| 3 | 20 | no | 1000 | 88 932 | 672.3 | 66 | 8 |
| 4 | 100 | no | 1000 | 88 635 | 668.7 | 66 | 41 |

Table S6: FRET dye and burst configuration parameters per $Mg^{2+}$ condition.

| $c_{\mathrm{Mg}}$ (mM) | $\tau_D$ (ns) | $\tau_A$ (ns) | dipole angle (°) | $Q_D$ | $Q_A$ | $\eta_D$ | $\eta_A$ |
|---|---|---|---|---|---|---|---|
| 0 | 1.33 | 1.28 | 18.81 | 0.65 | 0.51 | 0.88 | 1 |
| 10 | 1.40 | 1.28 | 16.59 | 0.43 | 0.35 | 0.88 | 1 |
| 100 | 1.67 | 1.42 | 17.45 | 0.46 | 0.37 | 0.88 | 1 |

Table S7: Anisotropy decay fit parameters (global model) from experiment and simulation. Global rotational correlation times $\tau_g$ and the derived rotational hydrodynamic radii $R_{\mathrm{h,rot}}$ were obtained from the water viscosity at 294.15 K ($\eta = 0.978$ mPa s, experiment) and 298 K ($\eta = 0.893$ mPa s, simulation). Uncertainties are the standard errors of the fit.

| source | dye | $c_{\mathrm{Mg}}$ (mM) | $\tau_g$ (ns) | $\tau_l$ (ns) | $r_\infty$ | $r_0$ | $\chi$ | $\theta_0$ (°) | $R_{\mathrm{h,rot}}$ (nm) |
|---|---|---|---|---|---|---|---|---|---|
| experiment | Cy3 | 10 | $11.67 \pm 0.66$ | 0.988 | 0.254 | 0.347 | 0.732 | 25.8 | $2.26 \pm 0.04$ |
| experiment | sCy5 | 10 | $14.29 \pm 1.57$ | 0.677 | 0.112 | 0.355 | 0.316 | 47.8 | $2.42 \pm 0.09$ |
| experiment | Cy3 | 100 | $17.58 \pm 0.67$ | 0.704 | 0.284 | 0.344 | 0.825 | 20.3 | $2.59 \pm 0.03$ |
| experiment | sCy5 | 100 | $27.08 \pm 2.71$ | 0.622 | 0.196 | 0.348 | 0.564 | 34.6 | $2.99 \pm 0.10$ |
| sim. restraint | Cy3 | 10 | $7.72 \pm 0.15$ | 0.562 | 0.270 | 0.334 | 0.809 | 21.4 | $2.04 \pm 0.01$ |
| sim. restraint | sCy5 | 10 | $10.45 \pm 0.21$ | 0.224 | 0.328 | 0.348 | 0.942 | 11.4 | $2.26 \pm 0.02$ |
| sim. restraint | Cy3 | 100 | $9.36 \pm 0.12$ | 0.288 | 0.312 | 0.337 | 0.925 | 12.9 | $2.18 \pm 0.01$ |
| sim. restraint | sCy5 | 100 | $9.47 \pm 0.15$ | 0.265 | 0.320 | 0.342 | 0.935 | 12.0 | $2.18 \pm 0.01$ |
| sim. unrestraint | Cy3 | 10 | $10.65 \pm 0.27$ | 0.392 | 0.252 | 0.335 | 0.752 | 24.7 | $2.27 \pm 0.02$ |
| sim. unrestraint | sCy5 | 10 | $11.08 \pm 0.22$ | 0.322 | 0.300 | 0.344 | 0.872 | 17.2 | $2.30 \pm 0.02$ |
| sim. unrestraint | Cy3 | 100 | $8.28 \pm 0.24$ | 0.557 | 0.180 | 0.329 | 0.547 | 35.4 | $2.09 \pm 0.02$ |
| sim. unrestraint | sCy5 | 100 | $10.51 \pm 0.18$ | 0.235 | 0.309 | 0.342 | 0.904 | 14.8 | $2.26 \pm 0.01$ |

Table S8: Mean burst FRET efficiency (explicit dyes and mACV), $E_{DA}$ without burst sampling under the isotropic $\kappa^2 = 2/3$ assumption, mean dye $R_{DA}$ and AP distance and mean $\kappa^2$ per $Mg^{2+}$ condition and structural state.

| $c_{\mathrm{Mg}}$ (mM) | state | $\overline{E}_{\mathrm{explicit\,dyes}}$ | $\overline{E}_{\mathrm{DA\,explicit\,dyes}}$ | $\overline{E}_{\mathrm{mACV}}$ | $\overline{E}_{\mathrm{DA\,mACV}}$ | $\overline{R}_{DA}$ (nm) | $\overline{AP}$ (nm) | $\overline{\kappa^2}$ |
|---|---|---|---|---|---|---|---|---|
| 0 | RNAComposer | – | – | 0.33 ± 0.05 | – | – | – | – |
| 10 | restraint | 0.90 ± 0.03 | 0.89 ± 0.05 | 0.92 ± 0.03 | 0.92 ± 0.02 | 4.21 ± 0.39 | 3.78 ± 0.18 | 0.438 |
| 10 | unrestraint | 0.67 ± 0.05 | 0.66 ± 0.08 | 0.85 ± 0.04 | 0.84 ± 0.04 | 5.44 ± 0.36 | 4.39 ± 0.24 | 0.297 |
| 100 | restraint | 0.91 ± 0.03 | 0.91 ± 0.04 | 0.93 ± 0.03 | 0.92 ± 0.02 | 4.22 ± 0.35 | 3.79 ± 0.24 | 0.375 |
| 100 | unrestraint | 0.76 ± 0.05 | 0.75 ± 0.11 | 0.91 ± 0.03 | 0.89 ± 0.04 | 5.11 ± 0.58 | 4.05 ± 0.30 | 0.412 |
